# Reduced amino acid substitution matrices find traces of ancient coding alphabets in modern day proteins

**DOI:** 10.1101/2025.02.19.639152

**Authors:** Jordan Douglas, Remco Bouckaert, Charles W. Carter, Peter R. Wills

## Abstract

All known living systems make proteins from the same twenty canonically-coded amino acids, but this was not always the case. Early genetic coding systems likely operated with a restricted pool of amino acid types and limited means to distinguish between them. Despite this, amino acid substitution models like LG and WAG all assume a constant coding alphabet over time. That makes them especially inappropriate for the aminoacyl-tRNA synthetases (aaRS) - the enzymes that govern translation. To address this limitation, we created a class of substitution models that accounts for evolutionary changes in the coding alphabet size by defining the transition from nineteen states in a past epoch to twenty now. We use a Bayesian phylogenetic framework to improve phylogeny estimation and testing of this two-alphabet hypothesis. The hypothesis was strongly rejected by datasets composed exclusively of “young” eukaryotic proteins. It was generally supported by “old” (aaRS and non-aaRS) proteins whose origins date from before the last universal common ancestor. Standard methods overestimate the divergence ages of proteins that originated under reduced coding alphabets in both simulated and aaRS alignments. The new model reduces this bias substantially. Our findings support the late incorporation of tryptophan into the genetic code (relative to tyrosine) and suggest that isoleucine and valine were once coded interchangeably, forming protein quasispecies. This work provides a robust, seamless framework for reconstructing phylogenies from ancient protein datasets and offers further insights into the dawn of molecular biology.

## 1 Introduction

> “*In the places I go there are things that I see*
>
> *That I never could spell if I stopped with the Z*
>
> *I’m telling you this ’cause you’re one of my friends*
>
> *My alphabet starts where your alphabet ends!*”
>
> - Dr. Seuss

While there are over five hundred types of amino acids in nature (Flissi et al., 2020), cellular systems employ just twenty-two (Atkins and Gesteland, 2002) to assemble proteins through genetic coding in a high-fidelity and nearly-deterministic fashion. What then was the nature of this coding table when life on Earth was first getting started? We argue that these codes would be both archaic and chaotic; using what little pool of amino acids was available to a low-fidelity, stochastic process that produced “statistical proteins” (Woese, 1965, 2004), or protein “quasispecies” (Wills et al., 2015), in a seemingly random fashion. After a series of expansions of the coding alphabet, refinements of the catalytic machinery, and optimisations of codon assignments (Koonin and Novozhilov, 2009), life eventually grew so complex that any further augmentation of the genetic code would become incremental and rare, giving the appearance of a “frozen accident” (Crick, 1968; Ribas de Pouplana et al., 2017; Kondratyeva et al., 2022).

Many lines of evidence paint a similar picture of the composition of the earliest coding alphabets; by placing the small, the branched-chain, and the acidic side chains among the first to appear; and the basic, the aromatic, and the amides among the last. As first demonstrated by the 1950’s Miller-Urey experiments (Miller, 1953; Johnson et al., 2008; Parker et al., 2011), some of the smaller amino acids – including alanine and aspartate – might have been produced in relative abundance under prebiotic conditions; while the larger and more reactive would appear later in protocellular history. Many of the supposedly “modern” amino acids have since become instrumental to protein function; for instance, histidine, the amino acid most commonly involved in catalysis (Ribeiro et al., 2020) is perhaps one of the youngest; and the latecoming positively charged residues (lysine and arginine) are now instrumental for facilitating protein-RNA binding (Ellis et al., 2007). Although there are some discrepancies in the order of amino acid appearance inferred from prior studies, the alphabet-expansion hypothesis is well-supported by the metabolic coevolutionary theory (Wong, 1975, 2005; Takénaka and Moras, 2020), the biophysics of protein folding (Makarov et al., 2023), phylogenetic reconstruction of ancient protein sequences (Brooks et al., 2002; Douglas et al., 2024a; Wehbi et al., 2024), and a statistical consensus model based on combined evidence from studies covering a range of amino acid properties (Trifonov, 2000). Furthermore, the intruder theory places the non-proteinogenic ornithine within the earliest of coding alphabets (Jukes, 1973); a code that was likely selected against due to the diminished foldability of proteins that contain ornithine (Makarov et al., 2023), and the instability of ornithinyl-tRNA (Weber and Miller, 1981; Hendrickson et al., 2021).

Any explanation of genetic code evolution must be considered in light of the aminoacyl-tRNA synthetases – the enzymes that attach amino acids to their cognate tRNA during codon-directed protein synthesis (**Fig. 1**), thus affecting the translation itself.

**Fig. 1:**
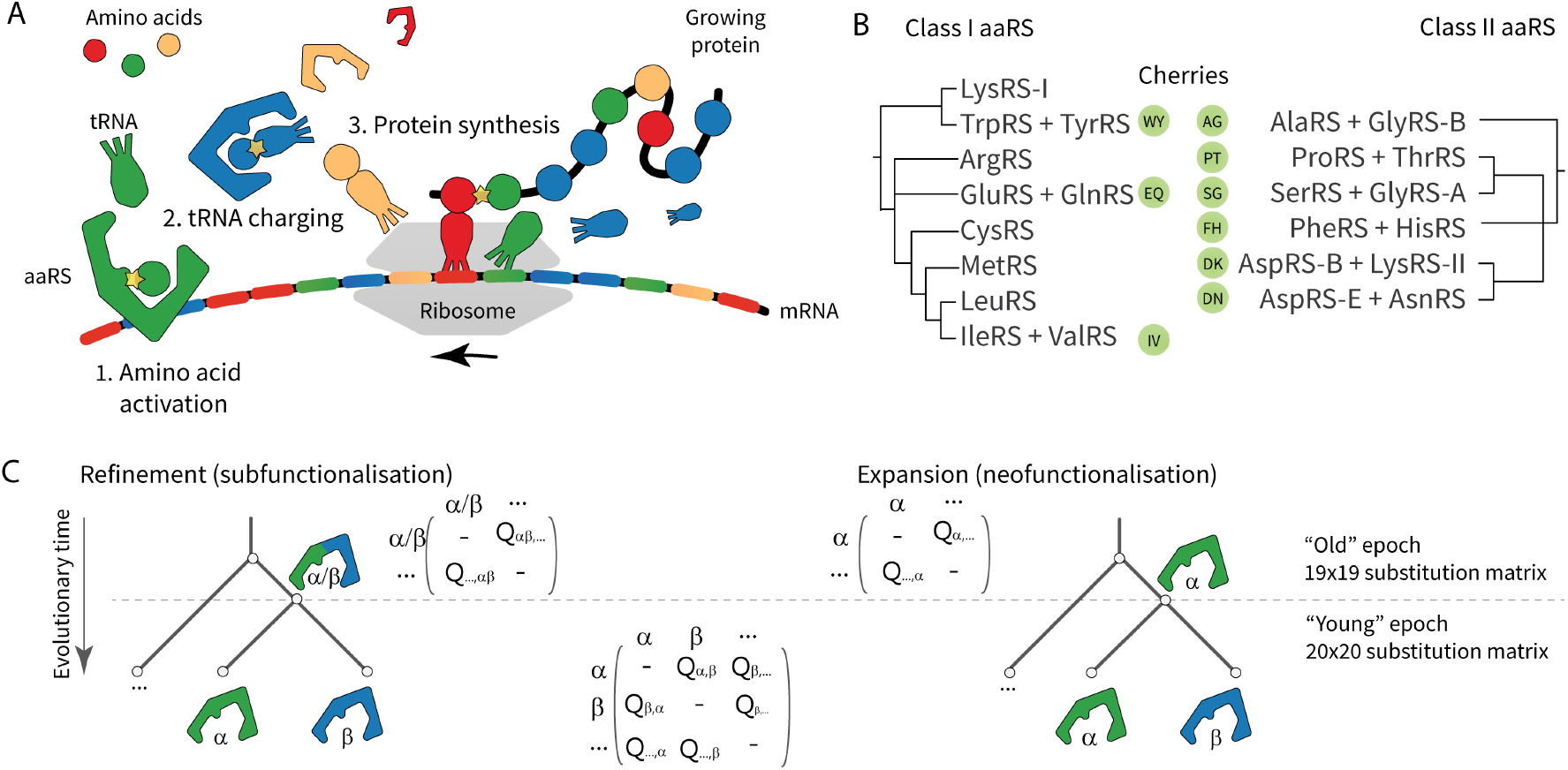
A: The aaRS attach amino acids to their cognate tRNA in ribosomal-directed protein synthesis. B: Cladogram of Class I and II aaRS based on Douglas et al. (2024c), with the three Class I and five Class II cherries indicated. Note that AspRS and GlyRS exist in distinct bacterial-like (B) and archaeal- or eukaryote-like (A and E) forms. C: Two possible hypotheses concerning the ancestral function of two closely related aaRS – *α*-RS and *β*-RS. Their ancestor might have recognised both amino acids, *α/β*-RS; or perhaps just one, say *α*-RS. Each hypothesis is associated with a different alphabet of amino acids and a different amino acid substitution model at the root of the tree.

### 1.1 Aminoacyl-tRNA Synthetases

The aminoacyl-tRNA synthetases (aaRS) have operated the genetic code in all living things since before the last universal common ancestor (LUCA). They do so by attaching amino acids to their cognate tRNA(s) in a two step reaction: first by activating the amino acid in the presence of ATP, and second by charging tRNA with the activated substrate (Gomez and Ibba, 2020). While the aaRS display many idiosyncrasies, in most instances each enzyme governs the expression of just a single type of amino acid, for example alanyl-tRNA synthetase (AlaRS) attaches alanine to the isotypes of tRNA^Ala^. aaRS catalytic domains fall into two seemingly unrelated evolutionary superfamilies - Class I and Class II (Eriani et al., 1990); which Rodin and Ohno (1995) hypothesise descended from the two complementary strands of a single bidirectional gene. This hypothesis has prompted biochemical investigations into the truncated, putative aaRS ancestors – known as “urzymes” – which have provided direct experimental evidence into how ancient alphabets might have operated (Li et al., 2011; Onodera et al., 2021; Carter Jr and Wills, 2021). Recent studies have demonstrated that Class I urzymes are better catalysts when apparently “modern” amino acids (histidine and lysine), now essential in the active site of full-length aaRS, are replaced with simpler alanine side chains (Tang et al., 2023, 2024).

Consider the ancestor of two aaRS species, *α*-RS and *β*-RS, in the canonical aaRS pathway. In one possibility, this ancestor was a low-specificity, promiscuous enzyme *α/β*-RS that was able to attach amino acids *α* and *β* to the same pool of tRNA species in a stochastic fashion. This form of non-deterministic genetic coding has been demonstrated in urzymes (Carter Jr and Wills, 2021; Patra et al., 2024), with sophisticated error correction systems having evolved to prevent this from happening too often in nature (Perona and Gruic-Sovulj, 2014). The differentiation of an ancestral gene into two aaRS with mutually exclusive functions would be a refinement in its specificity, or a *subfunctionalisation* event (**Fig. 1**; Rastogi and Liberles (2005)). In a second possibility, suppose the ancestor of *α*-RS and *β*-RS were an enzyme that rendered only one of the two, suppose *α*, in which case the bifurcation led to an expansion of the genetic code, to include *β*, that is, a *neofunctionalisation* event (Ohno, 2013). This has been proposed for TrpRS, which supposedly spawned off of TyrRS (Fournier and Alm, 2015). These two processes - refinement and expansion - would have left behind distinct footprints in modern day protein sequences (Fournier et al., 2011; Fournier and Alm, 2015).

Complementary to these two processes, we must also be mindful of alternative scenarios. First, through *parafunctionalisation*, two or more co-occurring aaRS species, some of which may now be extinct, might recognise the same amino acid (Fournier et al., 2011). Indeed many genomes possess multiple duplicates of the same aaRS species, often with distinct origins (Krahn et al., 2022; Radecki et al., 2024). Second, through *retrofunctionalisation*, older, low-specificity aaRS lineages might be salvaged and repurposed upon the emergence of novel amino acids (Douglas et al., 2024a). Many aaRS can recognise amino acids that are not commonly found in the cell (e.g., meta-tyrosine and canavanine, both of which are toxins (Klipcan et al., 2009; Hauth et al., 2023)). Therefore the distinction must be made between the “binding specificity“ of an aaRS, and its “biological function“ with respect to coding. Third, we must also keep in mind the scenario where an ancestral tRNA was recognised by several aaRS. For instance, an archaeal isoform of tRNA^Leu^ is acylated with both Leu (by LeuRS) and Met (by MetRS) in response to cooler temperatures (Schwartz and Pan, 2015),

We must also keep in mind, *pre-translational modification*, which provides an interesting departure from the canonical aaRS-tRNA pathway. Consider the aaRS for the two acid-amide pairs – GluRS/GlnRS (Class I) and AspRS/AsnRS (Class II). Since their respective diversifications happened post-LUCA, they occur in bacterial- and archaeal-like forms (Lamour et al., 1994; Hadd and Perona, 2014; Douglas et al., 2024c). The evolutionary histories of these aaRS are characterised by non-discriminating aaRS (AsxRS and GlxRS) that attach acidic residues (Asp and Glu) onto the amide tRNAs (tRNA^Asn^ and tRNA^Gln^), to be subsequently corrected by a second enzyme; an amidotransferase (Lamour et al., 1994; Becker and Kern, 1998; Becker et al., 2000). Therefore, the coding logic of these four amino acids is governed not only by the aaRS and tRNA, but the amidotransferases too. Pre-translational modification also enables the expression of cysteine in some archaea, using SepRS (Sauerwald et al., 2005); and the 21st amino acid selenocysteine in most organisms, using SerRS (Lee et al., 1989).

Placing these exceptions aside for the moment, the Class I and II aaRS phylogenies possess nine modern day *cherries* - defined as pairs of extant aaRS *α* and *β* that have distinct aminoacylation functions, and share an immediate ancestor with an unknown function; *α, β*, or *α/β* (Fig 1). These cherries are WY, IV, and EQ for Class I; and AG, SG, PT, FG, DN, and DK for Class II – many of which are composed of two amino acids with similar biochemical properties. Three cherries (AG, SG, and DK) contain an aaRS that has evolved along more than one evolutionary pathway (GlyRS and LysRS) and it remains unclear which aaRS, or perhaps both or neither, was respectively part of the predominant ancestral coding table (Valencia-Sánchez et al., 2016; Terada et al., 2002), while two other cherries (EQ and DN) are aaRS with amino acid substrates that are related through pre-translational modification. We will analyse each cherry to understand how the genetic code might have developed in early evolutionary history.

### 1.2 Challenging the phylogenetic fixed-alphabet assumption

Since Dayhoff et al. (1978), amino acid substitution models have become instrumental in characterising protein evolution (Whelan and Goldman, 2001; Le and Gascuel, 2008), with applications found in homology detection, sequence alignment, phylogenetics, and ancestral sequence reconstruction. These 20 × 20 matrices describe the exchangeability of amino acids as a result of their placement in the coding table and their biochemical properties, indicating the degree to which amino acids are interchanged pairwise over evolutionary timescales. However, these models are all based on the assumption of a constant 20 amino acid alphabet through evolutionary time. While this assumption might be reasonable for younger proteins (for example among animals or plants), it starts to become unjustifiable as the divergence times among taxa approach the origin of coding, around four billion years ago, and it becomes a logical absurdity for the description of earlier stages of aaRS evolution when the alphabet size was probably much smaller than 20.

In this study, we depart from the conventional fixed coding alphabet assumption and present a class of amino acid substitution models informed by the aaRS phylogeny. This is achieved by proposing two coding epochs – a 19 amino acid alphabet in the past, and a 20 state alphabet in the present, where the boundary between the two epochs is estimated – making an important step toward the ultimate goal of having one epoch per alphabet size. The hypothesis driving this model is that the aaRS phylogeny shaped the coding alphabet (Wills et al., 2015; Shore et al., 2020). Following the bioinformatic analyses of the IV and WY cherries by Fournier et al. (2011) and Fournier and Alm (2015), inference is herein performed under a hierarchical Bayesian phylogenetic framework, implemented in BEAST 2 (Bouckaert et al., 2019), which readily enables hypothesis testing, the quantification of uncertainty, and the modular incorporation of other biological processes such as branching models and molecular clocks.

## 2 Methods

Let 𝒯 be a rooted binary time-tree with *N* leaves (taxa) and *N* − 1 internal nodes. The height of a node describes its age since the present day (at height zero). Bayesian phylogenetic inference enables the estimation of 𝒯 and other parameters *θ* as probability distributions, as opposed to point estimates, based on observed data *D* (a multiple sequence alignment of *N* sequences and *L* sites). The posterior density is given by:

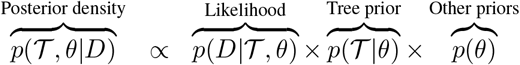

This distribution is sampled using Markov chain Monte Carlo (MCMC).

## 2.1 Substitution models with a fixed state space

The sequence likelihood *p*(*D*|𝒯, *θ*) is calculated using a dynamic programming algorithm that integrates over all possible ancestral sequences at the internal nodes of 𝒯 (Felsenstein, 1981). A transition probability matrix **P**(*τ*) is required, where entry *i, j* is the probability of transiting from state *i* to state *j*, and is a function of genetic distance *τ* between two points at different heights on the tree. It is assumed that each site evolves independently as a continuous time Markov process. Let **Q** be an *m* ×*m* instantaneous rate matrix, corresponding to the *m* = 20 standard amino acids (A, C, D, …, Y). **Q**_*i*,*j*_ describes the rate of substitution from *i* to *j*, for *i* ≠*j*:

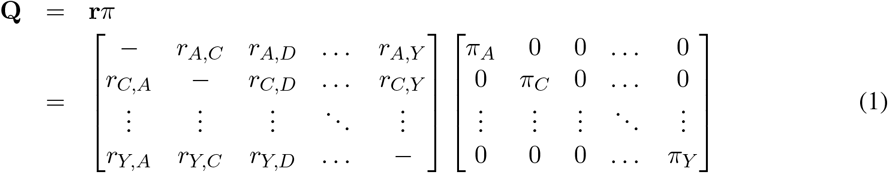

**r** is an exchangeability matrix whose elements reflect the ease with which one amino acid type can exchange with another over short evolutionary time frames, reflecting both random mutations in the encoding gene and the similarity between amino acids in the context of the corresponding folded protein. **r** is assumed to be symmetric, meaning that **r**_*i*,*j*_ = **r**_*j*,*i*_. The frequency of state *i* is represented by *π*_*i*_ = *π*_*i*,*i*_, such that Σ_*i*_ *π*_*i*_ = 1. The rows of the instantaneous rate matrix **Q** sum to zero by way of condition:

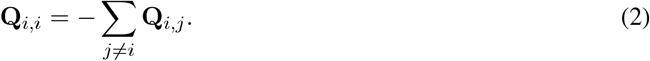

The entries in **Q** are normalised such that the mean outgoing rate is equal to 1 substitution per unit of time, giving *q*(**Q**).

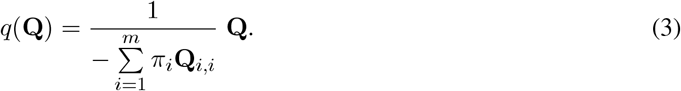

Finally, the transition probability matrix **P**(*τ*) is calculated from *q*(**Q**):

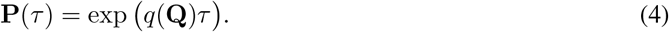

If a single lineage is associated with multiple substitution matrices **Q**_1_, **Q**_2_, **Q**_3_, …, their probability matrices can be multiplied together:

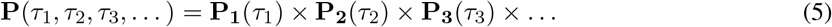

Empirical exchangeability matrices and frequency vectors are manifold, including BLOSUM62 (Henikoff and Henikoff, 1992), WAG (Whelan and Goldman, 2001), and VT (Müller and Vingron, 2000). Most such methods are time-reversible (meaning that **r** is a symmetric matrix), however there has been recent work on the estimation of asymmetric exchangeability matrices in which the reversibility assumption is relaxed (Dang et al., 2022). In a typical phylogenetic analysis, **r** is fixed at its empirical estimate, and *π* may be fixed or estimated as a parameter of the model under consideration. Here, however, we will be estimating both **r** and *π* during MCMC to avoid a multimodal posterior distribution that can result from model averaging (Bouckaert, 2020).

### 2.2 Substitution models with a two-epoch state space

Let *α* and *β* be two amino acids corresponding to two families of aaRS; *α*-RS and *β*-RS. The members of each family are monophyletic and the two families are monophyletic with each other, meaning they form a cherry on the aaRS family tree, with age *t*_*α*,*β*_. <u>**Example 1:**</u> *α* might be tryptophan (Trp) and *β* tyrosine (Tyr); where TrpRS and TyrRS are known to be monophyletic across the tree of life (Fournier and Alm, 2015; Douglas et al., 2024a).

Let resub(*α, β*) denote a **refinement-expansion substitution** model with respect to amino acids *α* and *β*. In this substitution model, we will relax the fixed-alphabet assumption by allowing for *two* coding epochs *E*_1_ and *E*_2_, which proceed backwards in time. The young epoch *E*_1_ = [0, *t*_*e*_] contains *m* = 20 amino acids, including *α* and *β*, and an amino acid substitution process described by exchangeability matrix **r** and amino acid frequencies *π*. The age *t*_*e*_ of this epoch is assumed to be close to the age of bifurcation *t*_*α*,*β*_ among these two aaRS, for example using a Laplace distribution:

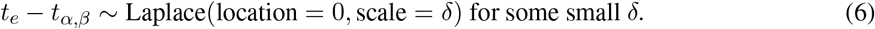

The old epoch is defined by *E*_2_ = (*t*_*e*_, *t*_*h*_], where *t*_*h*_ is the age of the root of . This epoch describes an earlier period in which there were effectively *m* = 19 coded amino acids, and this reduced alphabet is governed by the lower-dimension matrices **Q**^′^, **P**^′^, **r**^′^ and *π*^′^. Let *γ*_0_ ∈ {*α, β*} be the index of the state that was discarded from the old epoch, and let *γ*_1_ ∈ {*α, β*} be the index that is included, such that *γ*_0_ ≠ *γ*_1_.

There are three competing scenarios that can describe the composition of the reduced alphabet in *E*_2_. The model indicator 𝕀_*s*_ ∈ {0, 1, 2, 3}, which is estimated, describes the outcome according to the following four cases.

#### Case 0

If 𝕀_*s*_ = 0, both epochs share a common substitution process with *m* = 20 states. This is the null hypothesis where *α* and *β* persist in both epochs.

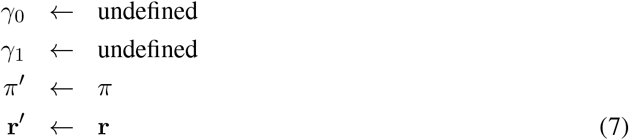

#### Case 1

If 𝕀_*s*_ = 1, the ancestor of *α*-RS and *β*-RS was a low-specificity enzyme *α/β*-RS unable to discriminate between amino acids *α* and *β*. The gene duplication event that spawned the two children led to a *refinement* of specificity (that is, a subfunctionalisation event). In this case, *α* and *β* are merged into *α*, and *β* is removed. This merged state is assumed to be at equilibrium between its sub-states. This equilibrium state is denoted as *α/β*.

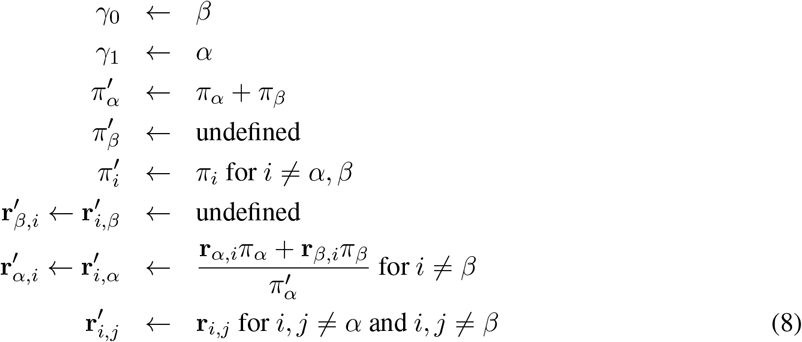

#### Case 2

If 𝕀_*s*_ = 2, the ancestor was *α*-RS, and thus *β*-RS represents a functional innovation, or an *expansion* of the coding alphabet (that is, a neofunctionalisation event). In this case, *β* is removed from the state space and the remaining frequencies are normalised so they sum to 1:

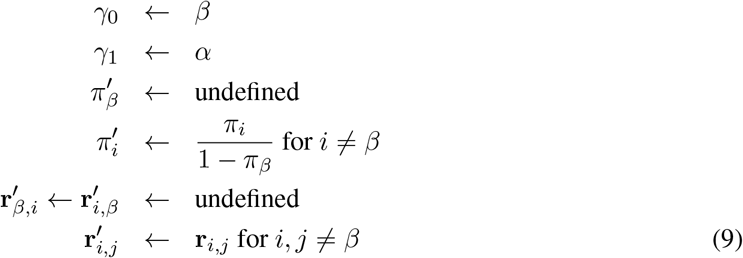

#### Case 3

If 𝕀_*s*_ = 3, then *α* is removed from the state space (same pattern as Case 2 but with *α* and *β* transposed).

When the null hypothesis is rejected (𝕀_*s*_ > 0), all lineages in *E*_1_ use the standard 20 × 20 substitution matrix, and those in *E*_2_ use the 19 × 19 form. Special attention is required for the case where a lineage crosses the epoch border. In this scenario, a 19 × 20 transport matrix 𝕋 transfers probability into the newly generated state. Suppose that a lineage crosses *E*_1_ with *τ*_1_ units of evolutionary distance, and *E*_2_ with *τ*_2_ units. Then, the Markov transition matrix of this lineage is:

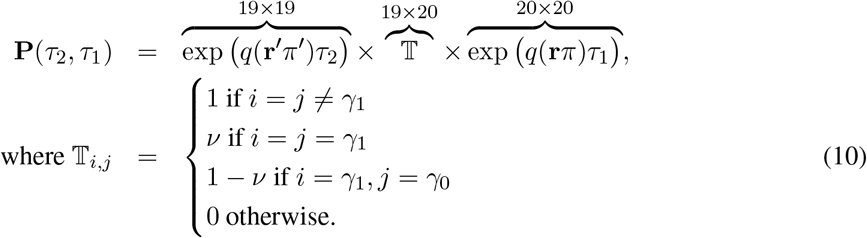

At this epoch boundary, it is assumed that each ancestral character *γ*_1_ remains in the same state with probability ν, or instantaneously transits to *γ*_0_ with probability 1 − ν. Each row in this matrix has a probability sum of 1.

<u>**Example 2:**</u> Suppose that *α* = W (tryptophan), *β* = Y (tyrosine), and 𝕀_*s*_ = 1. In this case, their ancestral aaRS held dual specificity. In the reduced matrix, the Y row and column have been removed (*γ*_0_ = Y), and the W row and column have been retained (*γ*_1_ = W), with the row renamed to W/Y in the matrix below:

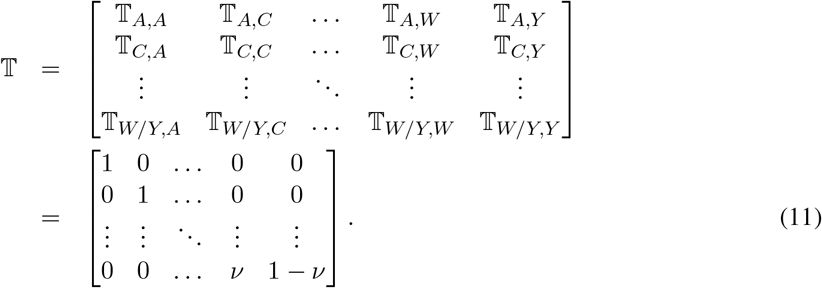

<u>**Example 3:**</u> Let us consider a fully worked toy example with *m* = 3 states denoted by x, W, Y in that ordering; where W and Y are the two characters we consider merging. This example is illustrated in Fig. 2. During *E*_1_, the **Q** matrix takes the form:

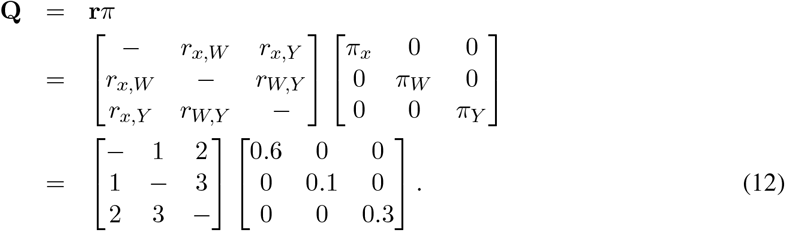

**Fig. 2:**
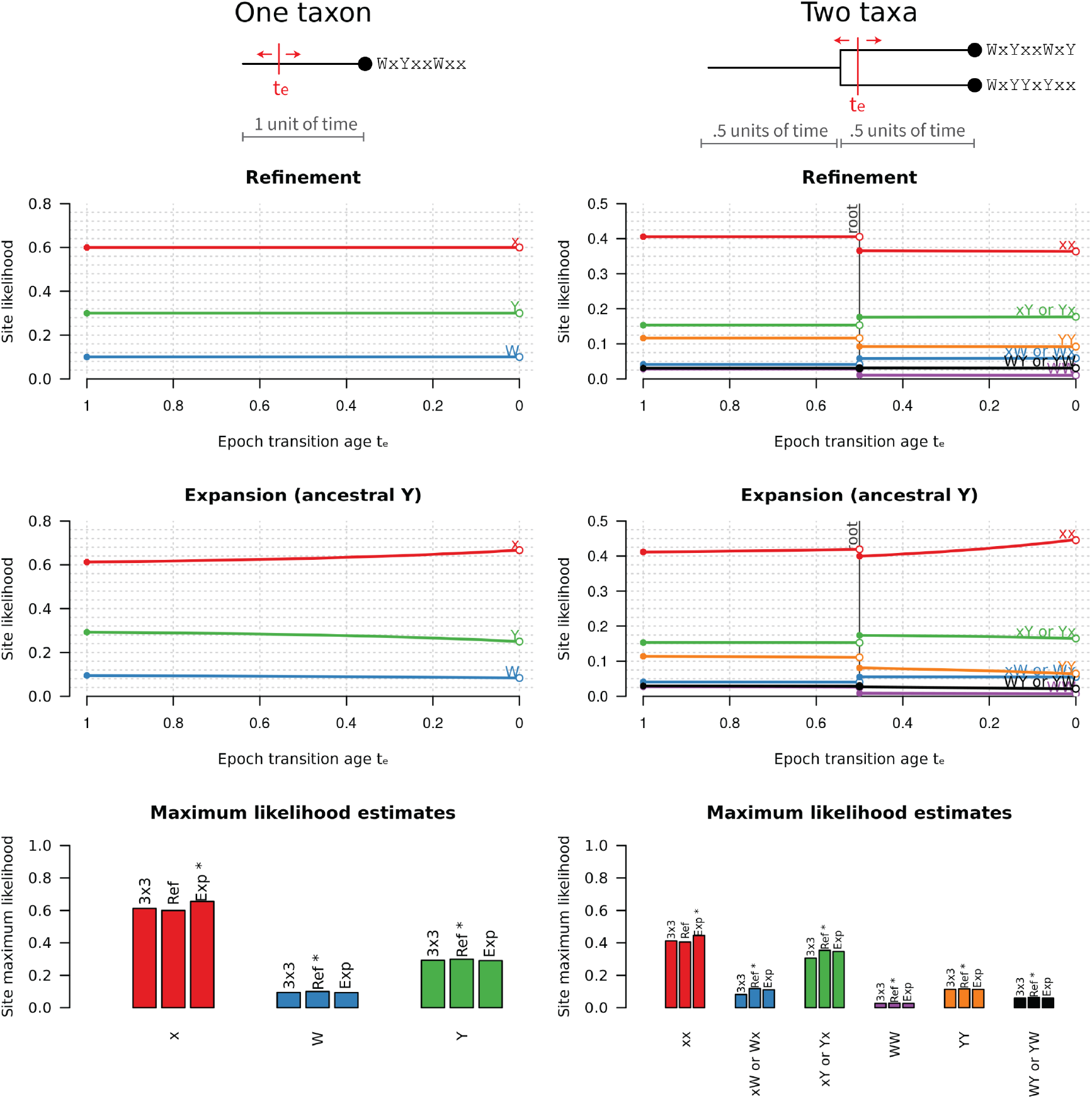
Site likelihoods under the resub model. In this toy example, we consider an alphabet of size *m* = 3 with states W, Y, and x; where W and Y are the two characters we consider merging. We explore a one-taxon tree with origin height 1 (left column) and a two-taxon tree with origin height 1 and root height 0.5 (right column). In the case of a one taxon tree, there are just 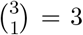 possible site patterns that any given column in the multiple sequence alignment can adopt - x, W, and Y. In contrast, a two taxon tree has 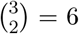 possible site patterns if we ignore taxon ordering - xx, xW (or Wx), xY (or Yx), WW, YY, and WY (or YW). These patterns have distinct likelihoods across varying *t*_*e*_, and across the refinement and expansion variants of resub. Note the discontinuity in likelihood for the two taxon tree. On the bottom row, the maximum likelihood (across varying *t*_*e*_) is reported for each combination of site pattern and substitution model (with the highest scoring model indicated by an asterisk). Altogether, this exercise demonstrates that multiple sequence alignments contain information that can subtly differentiate between varying resub models and the value of *t*_*e*_. 13

By normalising **Q** using the *q* function from Equation 3, and then rounding to three significant figures, we are left with:

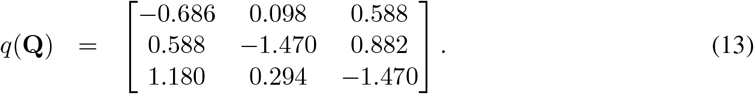

Suppose that *τ*_1_ = 0.5 substitutions per site have occurred. Then, the probability transition matrix during the young epoch *E*_1_ is:

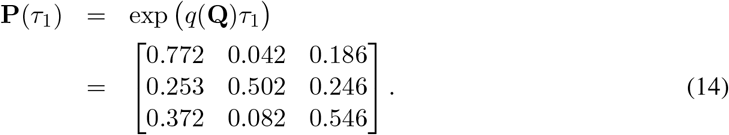

This matrix is directly used for calculating the tree likelihood using Felsenstein’s peeling algorithm (Felsenstein, 1981). Note that entry (*i, j*) is the probability of transitioning from row *i* to column *j*. Now, let us assume that the ancestral alphabet contains Y but not W (i.e., a case of expansion where 𝕀_*s*_ = 3). In this case, we will retain state index *γ*_1_ = Y and discard index *γ*_0_ = W from the reduced matrices. During epoch *E*_2_, the ancestral **Q**^′^ matrix is:

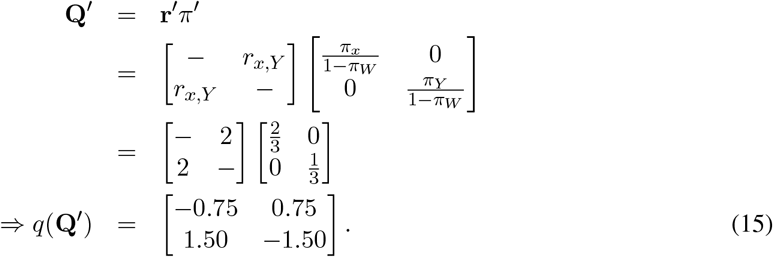

If *τ*_2_ = 0.5 substitutions per site have occurred, then the probability transition matrix during *E*_2_ is:

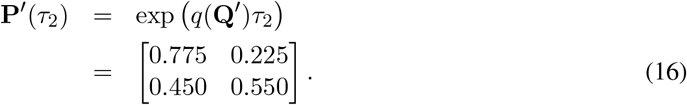

Lastly, suppose that a lineage spans *E*_1_ with *τ*_1_ = 0.5 units of distance, and *E*_2_ with *τ*_2_ = 0.5 units of distance. We will also assume that ν = 0.75 of all Y characters remain as Y while the remainder instantaneously transit to W at the epoch boundary. Then, the probability transition matrix across the epoch border is:

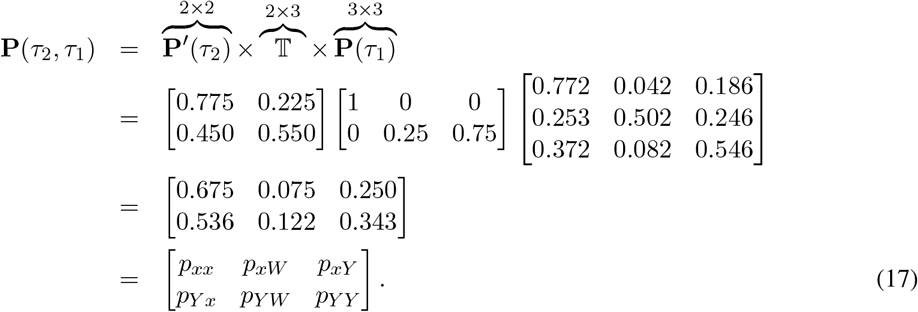

As shown in Fig. 2, the varying site patterns in a multiple sequence alignment do indeed give unique likelihoods under varying resub models, and varying values of the transition boundary *t*_*e*_.

### 2.3 Bayesian phylogenetic inference

Phylogenetic inference was performed using BEAST 2.7.5 (Bouckaert et al., 2019). Two independent MCMC chains were run on each configuration, ensuring that all reported parameters had a combined effective sample size over 200, as assessed by Tracer (Rambaut et al., 2018). Likelihood calculation times are accelerated using BEAGLE (Ayres et al., 2012). Posterior distributions of phylogenetic trees were summarised using the CCD-0 method (Berling et al., 2025). BEAST 2 XML files detailing the full evolutionary models are available online and the prior distributions are described in the Supporting Information, but in brief have the following components:

#### Substitution model

resub(*αβ*) for some *α* and *β* that form a cherry on the aaRS family tree. Each analysis is performed on a respective cherry *αβ*, which are shown in Fig. 1.

- Model indicator 𝕀_*s*_ for hypothesis testing.
- Amino acid frequency vector *π*.
- Proportion of sites that retain the old state at the epoch boundary transition ν.
- Age of boundary between the two epochs *t*_*e*_.
- Gamma site rate heterogeneity with four categories (Yang, 1994).
- Amino acid empirical exchangeability rates **r**.
- Each exchangeability rate has a corresponding boolean that specifies whether the rate is zero or non-zero, i.e., stochastic variable selection.

#### Clock model

In most analyses here, the gamma spike model was used to account for punctuated evolution (Douglas et al., 2024b). Built on top of the relaxed clock model (Drummond et al., 2006), this model assigns each lineage a spike of instantaneous change and a clock rate of gradual change. It was previously shown to be quite effective on aaRS phylogenies.

- Per-lineage relative rates.
- Log-normal distribution’s standard deviation of branch rates.
- Per-lineage spike size.
- Gamma distribution’s mean spike size.
- Gamma distribution’s shape of spike size.

#### Tree prior

Tree branching was assumed to follow a birth-death process (Nee et al., 1994), where per-lineage spike sizes are informed by the number of *stubs*, i.e., unobserved bifurcation events, under the gamma spike model (Douglas et al., 2024b).

- Lineage birth rate.
- Lineage reproduction number (equal to birth rate divided by death rate).

### 2.4 Divergence date priors

We used the divergence date estimates by Moody et al. (2024) in our Class I and II joint analysis. These dates were taken from the GMB model (partitioned + cross-bracing A). The lower and upper estimates were converted into LogNormal distributions with 95% interquartile ranges spanning this range, as shown in Table 1. These LogNormals then served as prior distributions in our own analyses. Cross-bracing (Shih and Matzke, 2013) was applied to ensure that the Class I and II tree roots occurred at around the same time. This was achieved by adding a Normal(µ = 0, *σ* = 0.01) prior on the difference between the two tree heights.

**Table 1:** Prior distributions used in divergence dating. The mean, lower, and upper were taken from Moody et al. (2024) - rounded to 3 significant figures in this table - and translated into LogNormal distributions with real-space mean M and standard deviation S, indicated above. All time units are in Ga.

| **Node** | **Mean** | **Lower** | **Upper** | **M** | **S** |
| --- | --- | --- | --- | --- | --- |
| Root | 4.50 | 4.45 | 4.52 | 4.49 | 0.00375 |
| Last universal common ancestor (LUCA) | 4.26 | 4.18 | 4.33 | 4.25 | 0.0094 |
| Last archaeal common ancestor (LACA) | 3.24 | 3.05 | 3.42 | 3.35 | 0.0114 |
| Last bacterial common ancestor (LBCA) | 3.24 | 3.16 | 3.34 | 3.25 | 0.0142 |
| Last mitochondrial common ancestor (LMCA) | 1.43 | 1.28 | 1.61 | 1.43 | 0.0589 |
| Last eukaryotic common ancestor (LECA) | 1.32 | 1.21 | 1.44 | 1.32 | 0.0443 |

## 3 Results

In the following sections, we demonstrate how the proposed two-alphabet model behaves on both simulated and empirical protein sequence data. The four hypotheses of the resub amino acid substitution model (I_*s*_ = 0, 1, 2, or 3) are compared using Bayesian model averaging (Wasserman, 2000; Hoeting et al., 1999). Much like the Akaike and Bayesian information criteria, Bayesian model averaging penalises overparameterised models, and is therefore expected to favour the simple null hypothesis (𝕀_*s*_ = 0) when the alternatives (𝕀_*s*_ > 0) and their two additional parameters, ν and *t*_*e*_, have little to offer. Competing hypotheses are compared using Bayes factors; with a threshold of 10 being considered as “strong” support in favour of one hypothesis over another (Kass and Raftery, 1995), corresponding to posterior support over 0.91 when two alternatives are equal *a priori*. We first validate and characterise this method through simulation studies, and then show the two-alphabet hypothesis is generally supported by Class I and II aaRS, but not by younger (non-aaRS) empirical datasets. Lastly, we apply resub to the Class I and II aaRS in a single joint analysis by linking the two phylogenies using divergence dates from across the tree of life. These experiments come with a high computational cost: for each empirical dataset, and for each cherry, two independent MCMC chains are run until they converge to the same posterior distribution; each chain taking several hours or days of computation time. Although one can never be certain that an MCMC chain has converged, comparing multiple independent chains and ensuring their effective sample sizes are large (over 200) is standard practice.

### 3.1 The true model is recovered from simulated data

We validated the correctness of this method using coverage simulation studies (Mendes et al., 2025). This was achieved by i) sampling parameters *θ* and a tree 𝒯 from specified probability distributions; ii) simulating a multiple sequence alignment *D* conditional on *θ* and 𝒯; iii) running MCMC on data *D* to obtain estimates 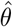 and 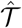; and lastly iv) comparing *θ* with 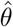 and 𝒯 with 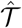. These four steps were iterated 100 times. If a model is correctly implemented, the known value of a term should lie in the 95% credible interval of its estimate in approximately 95% of all replicates. These experiments confirmed the usefulness and correctness of the method, whose estimates were well-correlated with the true values and had around 95% coverage, for varying taxon counts *N* and sequence lengths *L* (Fig. S1 and S2).

Second, we evaluated the method’s ability to identify the known substitution model. To do this, we simulated data under all four resub cases 𝕀_*s*_ ∈ {0, 1, 2, 3} and estimated I_*s*_ using MCMC, for varying taxon counts *N* and sequence lengths *L*. As shown in Fig. 3, the suitability of a two-epoch alphabet model is usually evaluated correctly even on small alignments (*N* = 20 and *L* = 100); and more so with increasing data volume. The true model (resub or no-resub) was usually identified with high support, and the incorrect model was favoured very rarely. However, as shown in Fig. S3, the method is much less powerful at discriminating between cases 𝕀_*s*_ = 1, 2, and 3. Distinguishing between refinement and expansion therefore requires a much larger volume of data than identifying the suitability of the two-epoch alphabet model in the first place. The observation that these substitution models can be discriminated at all is intuitive, since they receive different levels of support from varying site patterns (Fig. 2). Taken together, these results are reassuring; they confirm the method i) is statistically consistent, ii) is reliable at hypothesis testing even on small datasets, and iii) does not have a tendency to “overfit” to data generated by simpler evolutionary processes by using its two extra parameters.

**Fig. 3:**
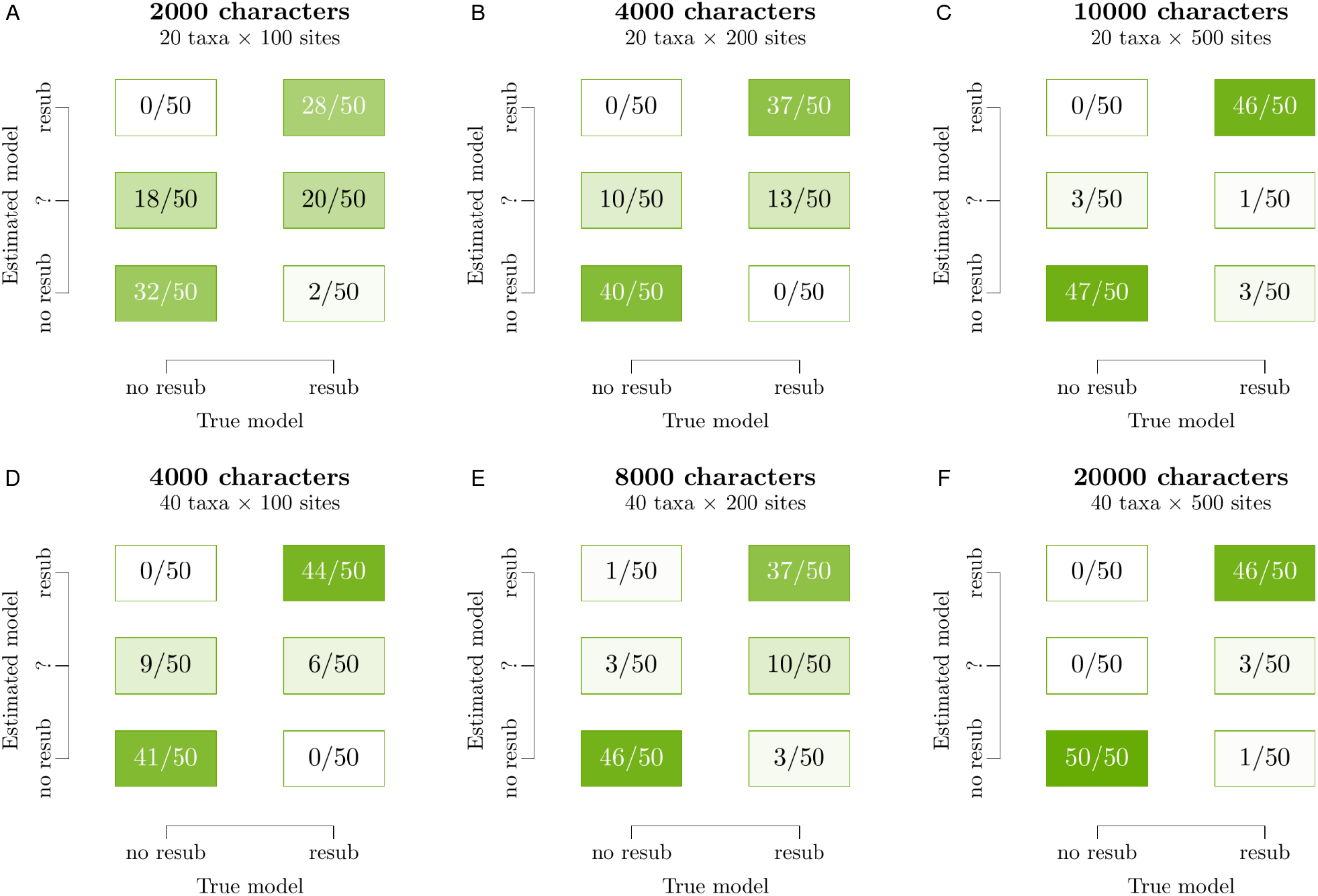
Identifying the suitability of resub on simulated data of varying sizes. For each panel in the figure, 100 datasets were simulated (50 under either substitution model) and an MCMC chain was run on each dataset. The inferred model was classified into “no resub” if *p*(𝕀_*s*_ = 0) > 0.91 or “resub” if *p*(𝕀_*s*_ ≠ 0) > 0.91, corresponding to a Bayes factor of 10. Replicates that yielded an intermediate posterior support are classified into “?”. These experiments show the known model can be recovered even on small alignments, and that the classifier becomes more confident in the correct answer with increasing volumes of data.

### 3.2 The two-alphabet hypothesis is supported by the aaRS but not by young proteins

We tested the two-alphabet hypothesis on three categories of empirical protein alignment: i) non-aaRS proteins found exclusively in eukaryotes and eukaryote-infecting-viruses (the putatively “young” proteins), ii) non-aaRS proteins with a likely bacterial, archaeal, or pre-LUCA origin (the “old” proteins), and iii) varying alignments of the Class I and II aaRS catalytic domains. Non-aaRS alignments were sampled from the expert-curated benchmark alignment database BAliBASE 3.0 (Thompson et al., 2005), such that each alignment had significant sequence divergence (with the tree height estimated as over 1.5 substitutions per site), and was not homologous with aaRS Class I or II catalytic domains. aaRS datasets were obtained from a curated database of aaRS alignments (Douglas et al., 2024c), and modified into three variants per Class, composed of i) the common elements of the catalytic domain, ii) urzyme only (i.e., a subset of the catalytic domain), and iii) urzyme with non-bacterial sequences removed. Thus, in total we sampled four young alignments and six old alignments from BAliBASE, and produced six variations of the aaRS catalytic domain. These sixteen multiple sequence alignments had a broad diversity of sequence and site counts (Fig. 4). They were screened against all nine putative cherries of the aaRS phylogeny (EQ, IV, WY, AG, SG, FH, DN, DK, and PT) as well as three negative controls – “fake” cherries that do not form clades on the aaRS phylogenies but instead occur across the two Classes (LS, CF, and NQ). As shown in Table S2 and S3, all twelve of these amino acid pairs were well-represented across the sites in the multiple sequence alignments, and as shown in Fig S4, there is a non-zero estimated rate of transition between each pair of these amino acids in the aaRS. All datasets were run under the same phylogenetic model, but with varying cherries; where the alphabet epoch transition *t*_*e*_ was estimated under the *a priori* assumption that it occurs near the root of the tree, without being strongly constrained to any particular ancestral node. In other words, we treated both types of data (aaRS and non-aaRS) in exactly the same manner.

**Fig. 4:**
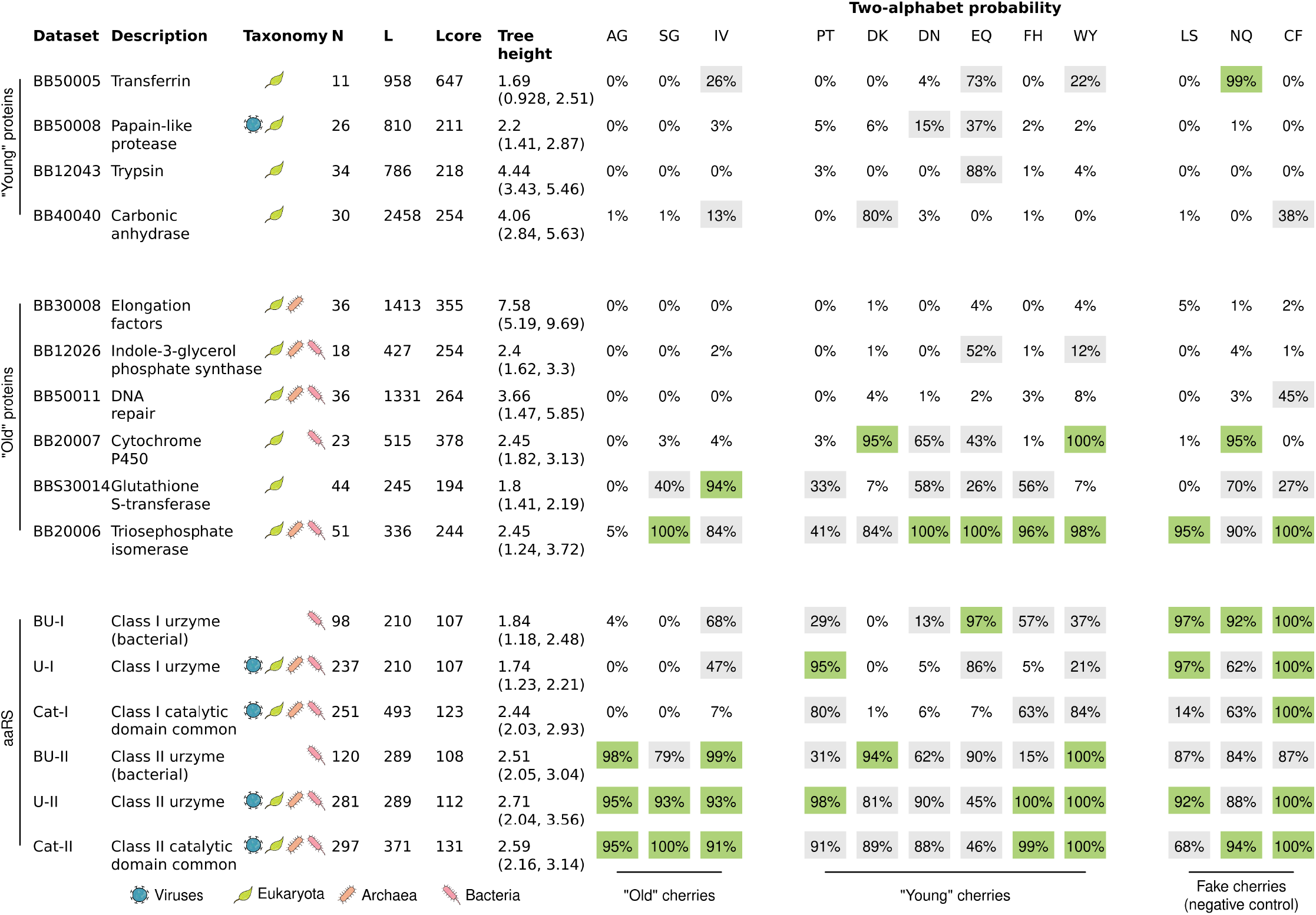
Testing the two-alphabet hypothesis on biological data: “young” eukaryotic/viral protein alignments, “old” proteins found beyond eukaryotes and viruses, and aaRS proteins. Columns: the name of the dataset, which is the BAliBASE (Thompson et al., 2005) identification number in the case of non-aaRS proteins, number of taxa *N*, number of sites *L*, number of sites that have < 25% gaps *L*_core_, estimated tree height (and 95% credible interval) in units of amino acid substitutions per site, and the posterior probability of the resub model being chosen for different cherries. The green squares represent strong support in favour of the two-alphabet hypothesis (with respect to a certain cherry), the white squares strong support against, and grey squares have intermediate support; at a Bayes factor threshold of 10. “Young” cherries are defined as those that contain at least one phase II amino acid according to Wong (2005). This experiment confirms that resub is often chosen on aaRS datasets, rejected on young non-aaRS proteins, and returns mixed results for old non-aaRS proteins; but it also returns some unexpected conclusions for the fake cherries.

This experiment was designed to verify several anticipated properties of the new method. First, we would expect the younger protein families to generally reject the resub model because they were most likely born under a contemporary coding table with 20 amino acids. However in some cases the model might still be supported for unrelated reasons; for example resub(WY) could be favoured even on a younger tree if several Y residues were to rapidly change across multiple lineages around the same point in time for whatever reason. Second, we would expect the Class I and II aaRS datasets to favour the new model with respect to each cherry, noting that we did not incorporate any prior information into the relationship between *t*_*e*_ and the aaRS ancestral nodes during this experiment. Third, some of the “old” protein datasets would favour the new model – particularly the aaRS bifurcation events that supposedly occurred later in time (e.g., WY, EQ, and DN) – while others reject it or have intermediate support. Lastly, the fake cherries (LS, CF, and NQ) should not be supported by any datasets, because they do not describe an aaRS functional bifurcation.

This experiment confirmed most of our hypotheses, further corroborating the reliability of the method, and also highlighting a major limitation (Fig. 4). Namely, resub was usually met with low support among the younger proteins and high support among the aaRS, while the older non-aaRS proteins gave mixed results. The Class II aaRS provided greater support for the two-alphabet model compared with Class I, despite the lack of prior information linking the alphabet transition boundary to cherry nodes in either case. Class II was met with greater cherry support with increasing dataset size; from bacterial urzyme, to urzyme, to catalytic domain common elements. Two of the non-aaRS datasets also provided strong support for resub: i) triosephosphate isomerase (TPI) – a glycolytic enzyme that was possibly found in LUCA (Moody et al., 2024), and ii) the glutathione S-transferase (GST) superfamily, which is likely of bacterial origin (da Fonseca et al., 2010) but unlikely to be found in LUCA (Moody et al., 2024) (note that there were no bacterial sequences in the sample, but multiple GST paralogs were included). Larger and more carefully assembled TPI and GST datasets would offer further clarity to these findings, which is beyond the scope of this study. Lastly, the negative control test failed, meaning that the fake cherries were often identified as real on the aaRS datasets, which may reflect a multitude of alphabet reduction events that cannot be readily disentangled without further information (such as aaRS bifurcation events). Interestingly, the fake cherries were still rejected by most of the non-aaRS datasets.

Taken together, this experiment i) provided reassurance that resub is describing a real biological process and is not just capturing random noise in the data, and ii) demonstrated that aaRS and other ancient protein sequences have retained useful signals about their primordial origins. It also underscores the need to incorporate further prior information if we are to extract richer information from the aaRS.

### 3.3 Class I and II aaRS evolved simultaneously

We evaluated the two-alphabet hypothesis on the Class I and II aaRS trees together. In order to coordinate the two phylogenies onto the same timescale, we used divergence date estimates from a recent tree of life study (Moody et al., 2024), presented in Table 1. These time calibrations were applied to the last universal (LUCA), bacterial (LBCA), archaeal (LACA), eukaryotic (LECA), and mitochondrial (LMCA) common ancestors across the aaRS, as well as the root of the two trees (near the origin of life). We note that the origin of life estimated by the previous study (4.49 Ga) is only marginally younger than the age of the planet (4.54 Ga; Dalrymple (2001)). The Class I and II tree root ages were further linked using cross-bracing (Shih and Matzke, 2013). One dated tree analysis was performed on each cherry, where the alphabet transition was assumed to affect both trees at the same point in time *t*_*e*_, and the timing of *t*_*e*_ was assumed to be around the same time as the corresponding aaRS bifurcation node on one of the two trees.

The catalytic domain alignments were generally met with greater support than the urzyme alignments, reflecting the availability of more data in the former case. In the case of the catalytic domain, these experiments provided strong support (>91%) in favour of the two-alphabet model for five cherries: WY, EQ, and IV for Class I; DN and SG for Class II. Whereas, the FH and PT cherries were met with intermediate support (51% and 10%), and the DK and AG cherries were strongly rejected (<1%). As a negative control, we also tested three fake cherries (LS, CF, and NQ), whose corresponding aaRS occur across the two Classes. Two of the three cherries were met with unexpectedly strong support (NQ and CF at over 98%), while LS was inconclusive. This finding suggests that other evolutionary processes might have been at play. Potential explanations may include the presence of tRNAs that were acylated by multiple aaRS, or the presence of multiple coding epochs. Further investigation is required to understand this unexpected result.

Very little clarity was offered as to the ordering in which the amino acids arrived - as demonstrated by the overlapping transition time posterior estimates (green curves in Fig. 6). The only information we can extract is that serine and glycine may have been differentiable in the coding table for much longer than the other pairs of amino acids considered. Further investigation is required to better understand the relative ordering between these alphabet expansion events e.g., a single analysis that combines the cherries together. The relationships between aaRS functional families according to resub are generally in line with our previous phylogenies (Douglas et al., 2024a,b).

### 3.4 Phylogenetic analysis is overestimating the age of life

Finally, we assessed the consequences of using the wrong substitution model to estimate root age. We hypothesised that application of a one-alphabet substitution model to data generated by multiple succeeding alphabets would result in an overestimation of root age. Our reasoning was that the fixed alphabet assumption cannot explain the myriad of amino acid substitutions that result from changing the genetic code, unless there is a longer period of time for those changes to occur independently. To test this hypothesis, we examined i) sequence data simulated under the two-alphabet model, and ii) the Class I and II aaRS catalytic domain trees, which also evolved from a reduced alphabet. In both cases, a subset of tree nodes were calibrated to specified dates, but the root was left uncalibrated. Then, we compared the root ages estimated under the one-alphabet and two-alphabet models.

To confirm that this bias is an expected property of the model, we simulated 50 alignments under the resub model with varying cherries, *L* = 200 amino acid sites, *N* = 40 extant taxa, and a variable number of non-extant taxa (ranging from 1 – 146 per tree) whose ages informed our calibrated clock model; detailed in Supporting Information Section 1.3. These trees were simulated under a birth-death-sampling process (Stadler, 2010). Then, we estimated the tree root height *t*_*h*_ and duration of the old epoch *t*_*h*_ − *t*_*e*_ using MCMC. This confirmed that the epoch duration *t*_*h*_ − *t*_*e*_ was consistently overestimated when the coding alphabet was fixed (𝕀_*s*_ = 0), in this case by an average of 68%, resulting in a 7% overestimation of the tree height *t*_*h*_ (Fig. 7). Unsurprisingly, the tree height was estimated more accurately when applying the resub model (𝕀_*s*_ > 0); with *t*_*h*_ − *t*_*e*_ being overestimated by only 17% and *t*_*h*_ by just 0.4%.

To confirm that this effect persists on empirical data as well as simulated, we reinterpreted the aaRS catalytic domains, but with weaker prior constraints placed on the two tree roots. As shown in Table 1, the root prior that we used in our earlier analysis was heavily constraining the age of the tree to be younger than the age of the Earth (4.54 Ga); by placing 95% of the prior probability between 4.46 and 4.52 Ga. This in turn resulted in our trees being just marginally younger than the Earth (Fig. 6). Therefore, in order to disentangle the effects of this prior from the signal coming from the aaRS data, we removed this stringent constraint, but retained all other divergence time calibrations (LUCA, LBCA, LACA, LMCA, and LECA). As shown in Fig. 7A, the new tree height prior distribution allowed the tree heights to adopt a much broader range of values, with a 95% credible interval of 4.46 – 5.70 Ga. Due to the large number of aaRS diversification events that took place before LUCA (at 4.25 Ga), this “prior distribution” has support for ages older than the Earth, but is much more permissive than our earlier one. Then, we compared aaRS root ages estimated under the default one-alphabet hypothesis (the Null model) with those of the favoured resub cherries (WY, IV, EQ, DN, and SG). This was performed under a relaxed clock model that allows the molecular clock rate to vary among lineages independently and from an identical distribution (Douglas et al., 2021), and a birth-death skyline model that allows the diversification rate to vary independently before and after LUCA (Stadler et al., 2013).

This experiment confirmed that resub gave younger tree ages than the Null model when applied to the aaRS (Fig. 7A and B). While all six estimates were significantly older than the age of the planet, the five cherries were in fact younger than the Null model: on average 12.2% younger for EQ, 12.1% for IV, 4.3% for SG, 3% for WY, and 0.8% for DN (each at *p* < 10^−12^ with a two-sided Kolmogorov-Smirnov test). If we are to assume that life did not originate extra-terrestrially, then the resub model therefore gave much more accurate time estimates. These age estimates were 28% closer to the age of the Earth for EQ and IV, 9.9% for SG, 6.9% for WY, and 1.8% for DN. A model with multiple coding epochs, rather than just two, is likely to yield even younger tree ages by combining these effects. The observation that some cherries have a much larger impact on tree height estimation than others can be partially explained by the frequencies of their amino acids in the aaRS alignment (Table S1). IV (i.e., I and V) has the greatest combined frequency (0.13), followed by SG (0.12), EQ (0.11), DN (0.075) and finally WY (0.068). It is intuitive that an instantaneous alphabet refinement or expansion event would have a greater impact on divergence time estimates when the two amino acids are more prevalent, for example I and V comprise 13% of the protein sequence on average and gave a 12% age reduction, while W and Y sit at just 7% and gave a 3% reduction.

Taken together, these results suggest that standard phylogenetic methods are overestimating the ages of proteins that evolved from a reduced alphabet, and therefore overestimating the age of life. This is an intuitive result; if one does not account for sudden, saltational changes in the proteome that result from coding table restructurings, they will overestimate the time taken to make those transitions. These rapid bursts of evolutionary change challenge the standard gradualistic assumption of clock-like molecular evolution (Pagel et al., 2006; Manceau et al., 2020; Douglas et al., 2024b). Standard one-alphabet methods also do not account for the fact that the older alphabet was smaller, and therefore has fewer states to mutate between. Our model here captures both of these phenomena and provides a partial correction for the bias.

## 4 Discussion

In this study, we challenged the standard phylogenetic assumption of a temporally constant amino acid coding-alphabet. This was achieved by developing a Bayesian phylogenetic method informed by the phylogeny of the aaRS; the enzymes that shaped the coding alphabet in early evolutionary history. The two aaRS phylogenies were respectively analysed in pairs of closely related aaRS, known as cherries (Fig. 1). We validated this approach using simulation studies, which confirmed the method’s effectiveness at testing the fixed-alphabet hypothesis (Fig. 3). Then, we demonstrated that the two-alphabet hypothesis is i) generally rejected by younger (non-aaRS) protein superfamilies that supposedly emerged after the coding table had matured; and ii) often supported by older (aaRS and non-aaRS) proteins (Fig. 4). The two-alphabet model met its strongest level of support when applied to the two aaRS Classes in a joint analysis, where the Class I and II aaRS phylogenies were temporally linked using divergence dating, and the alphabet transition heights were linked to aaRS bifurcation events. In this analysis, the divergence dates were predominantly informed by a recent tree of life analysis (Table 1); noting that alternate calibrations would result in alternate timelines (Fig. 5 and 6).

**Fig. 5:**
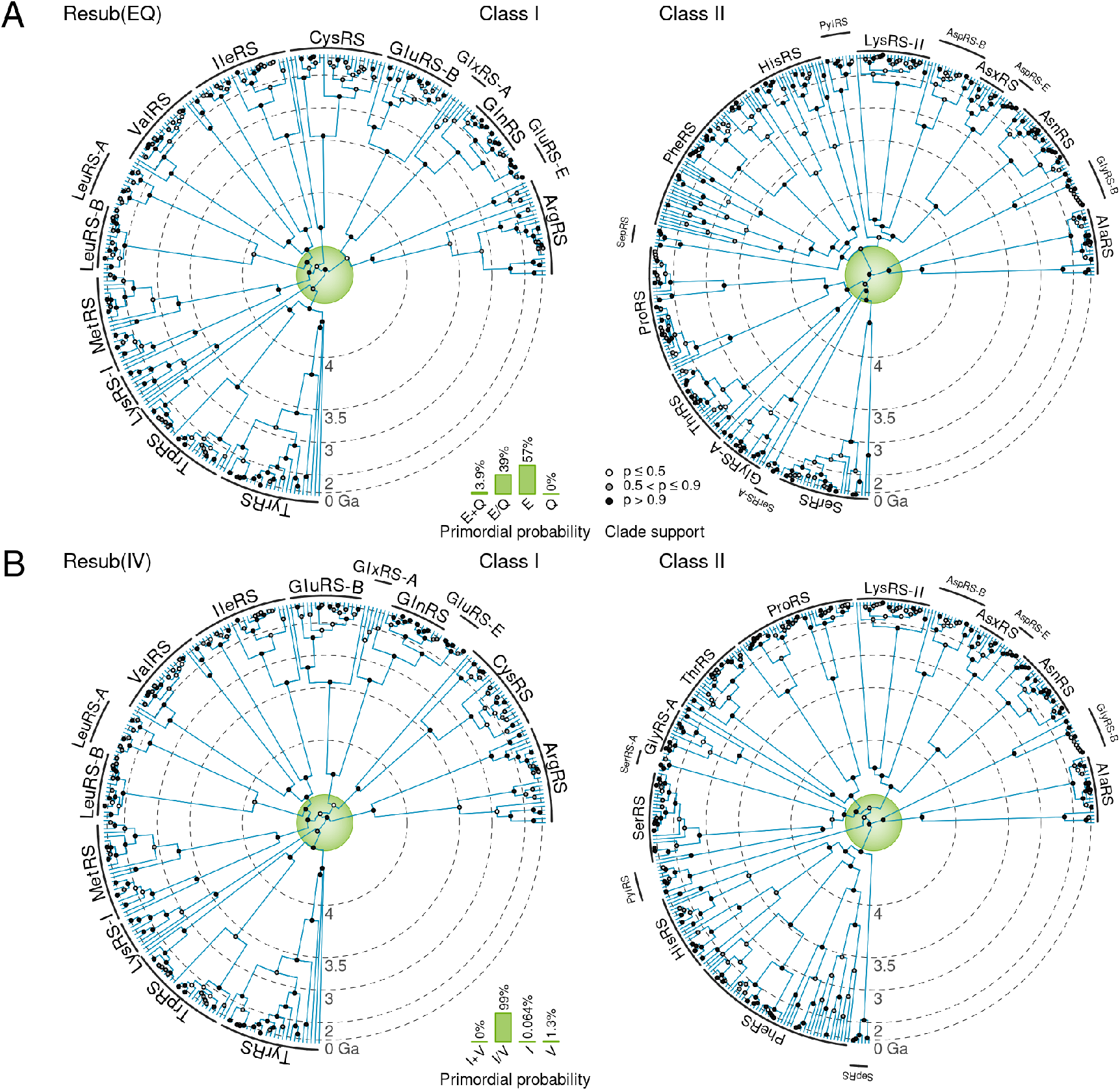
Class I and II catalytic domain phylogenies constructed using resub with A: the EQ cherry, and B: the IV cherry. In each case, the two trees were calibrated onto the same timescale using tree of life divergence date estimates. The inner green circle represents the old epoch *E*_2_ (with 19 amino acids), and the outer white region depicts the new epoch *E*_1_ (with 20 amino acids). Note that the timeline is on an exponential scale from the circumference to the centre, to highlight the fine structure within the old epoch. A: In this model, the 19 amino acid alphabet contains all 20 standard amino acids but with glutamate and glutamine either merged into a single state (with posterior probability 0.39), or with glutamate only (probability 0.57). B: The reduced alphabet is estimated to have isoleucine and valine merged into a single state (probability 0.99).

**Fig. 6:**
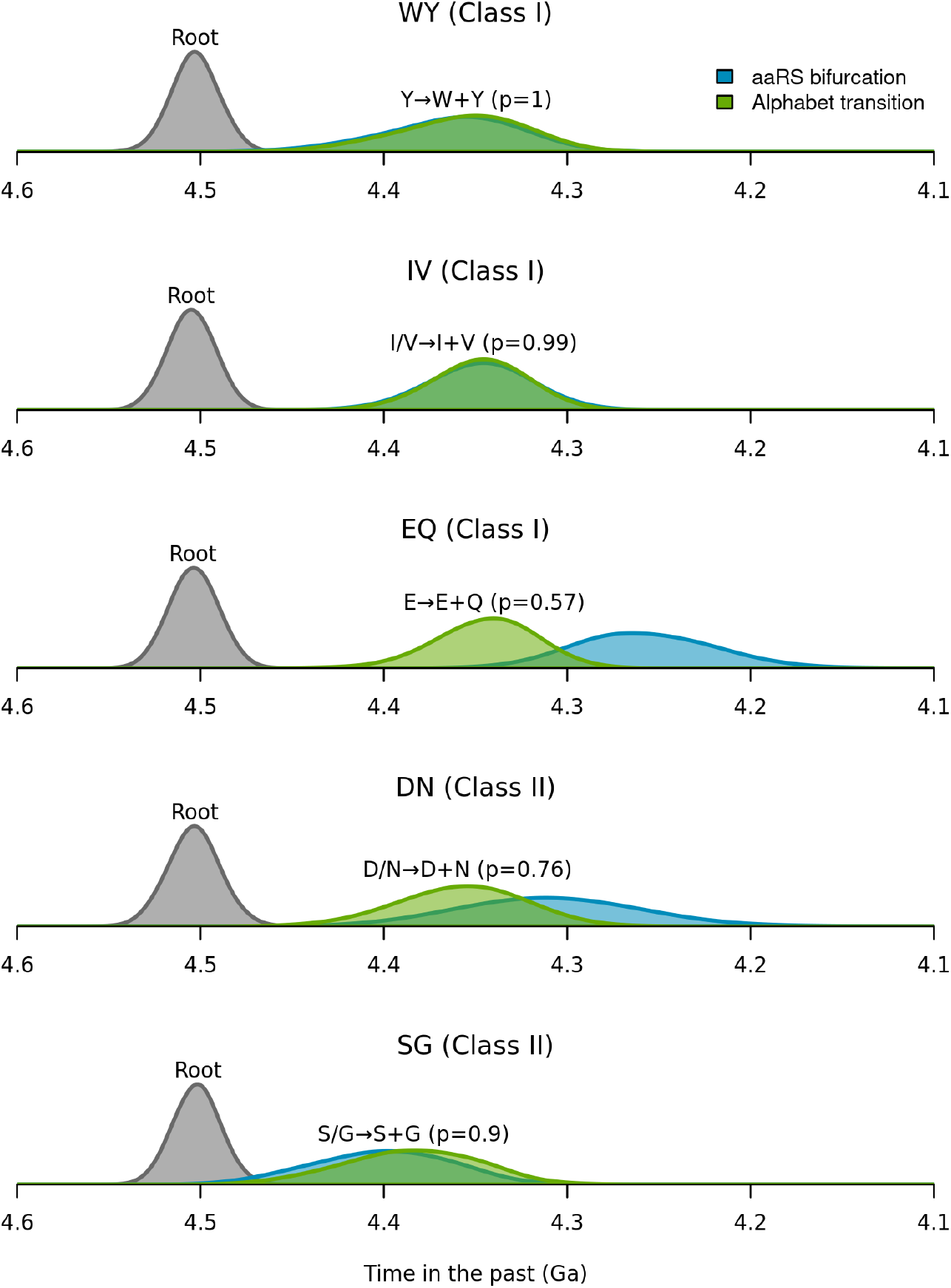
Estimated timeline of aaRS and code bifurcations. Each panel is a different aaRS cherry analysis that found high support (> 91%) for the two-alphabet hypothesis (Cat-I and II dataset). The y-axes are posterior probability densities of the age estimates. Notation: root – the average of the two Class tree roots; aaRS bifurcation – when the two aaRS are estimated to have diverged; Alphabet transition – when the reduced amino acid alphabet is estimated to have transitioned to the full one. We note that the dates shown here are predominantly informed by the calibrations from a previous study (Table 1); using alternative calibrations would yield a new timeline.

**Fig. 7:**
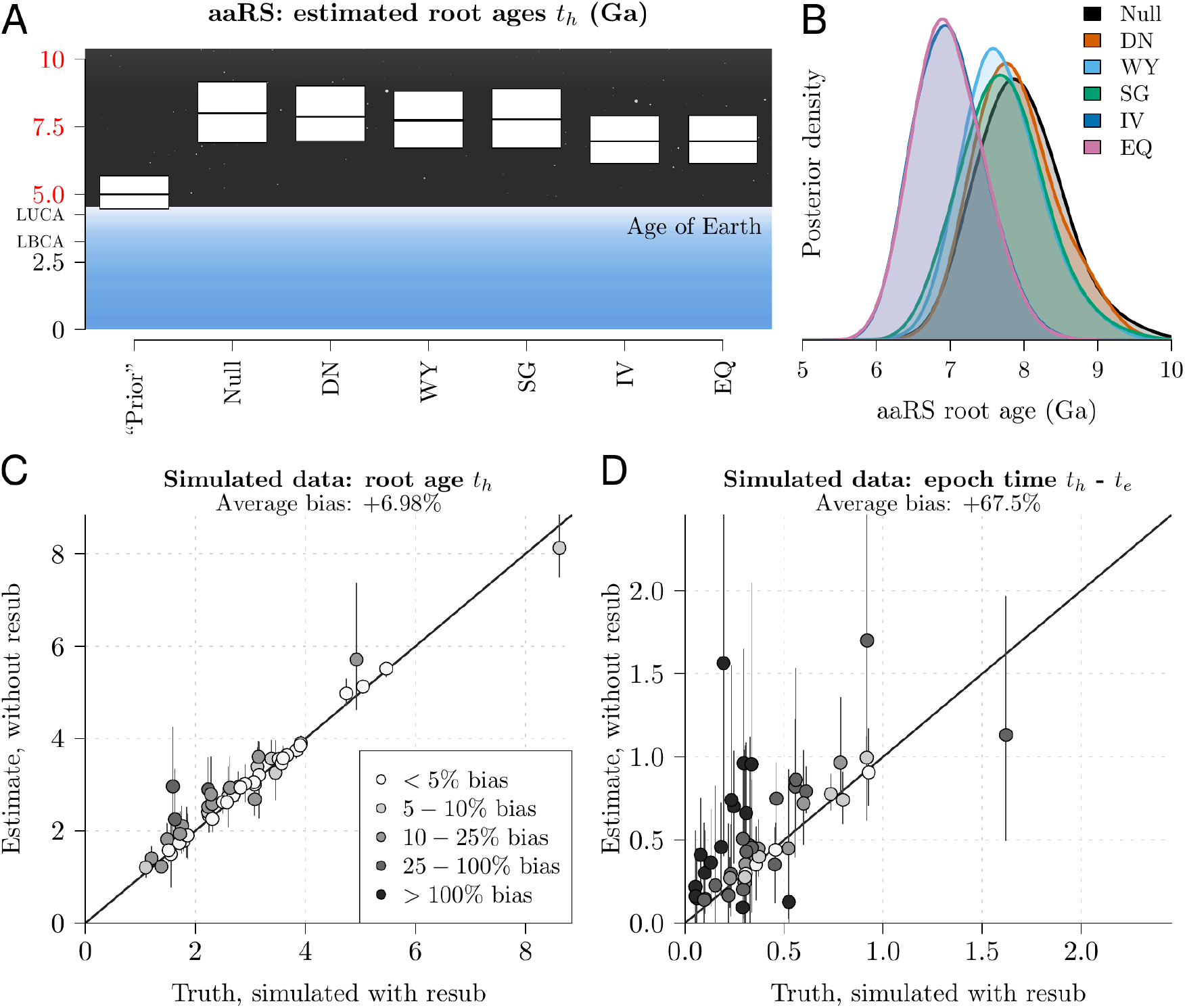
Overestimation of tree ages. A and B: The aaRS Class I and II tree ages were estimated using a standard 20 × 20 substitution model (Null), and with five different resub models. The white boxes in A depict means and 95% credible intervals, while the curves in B show the full distribution. Note that, in contrast to Fig. 5, here the tree roots were not constrained to be younger than the Earth, as shown by the comparatively broad prior distribution in A. Therefore all age estimates above ≈4.25 Ga are model-based extrapolation (red axis colours). According to these phylogenetic methods, the aaRS are significantly older than the planet; reflecting the absurdity in an assumption of the same clock-like evolutionary process applying throughout all of life’s history. However, the resub model partially corrects this overestimation. C and D: Using data simulated under a two-alphabet model, we estimated the tree height *t*_*h*_ and duration of the old epoch *t*_*h*_ − *t*_*e*_ under an assumption of a constant coding alphabet. This experiment shows that the standard one-alphabet amino acid model significantly overestimated the duration of the old epoch and therefore overestimated the age of the tree. Mean estimates are depicted by points, 95% credible intervals by vertical lines, and the reported biases are relative to the true known value.

### 4.1 Finding traces of ancestral coding alphabets in modern day proteins

Remarkably, despite undergoing four billions years of evolutionary change, today’s aaRS genes have preserved enough information to peer deep into life’s history, to a time when they were translated using alphabets of < 20 amino acids (Fournier and Alm, 2015; Shore et al., 2020). By leveraging this information, our experiments corroborated several existing hypotheses; further validating this approach. First, our findings confirmed that Tyr most likely predated Trp in the coding alphabet. This is an intuitive finding given the longer metabolic pathway needed to produce the bulkier Trp (Takénaka and Moras, 2020), and is in line with previous sequence analyses (Fournier and Alm, 2015).

Second, our results suggest that Ile and Val were once used interchangeably, with their coding likely implemented by a promiscuous enzyme that would stochastically attach either amino acid to the same anticodon’s tRNA (Fig. 5); thereby producing protein quasispecies (Wills et al., 2015). This result is consistent with LeuRS urzyme experimental studies (Carter Jr and Wills, 2021) and is also intuitive, given the structural similarities of Ile and Val, and the extensive error-correction machinery that has since evolved to ensure IleRS and ValRS can expel each other’s substrate (Fersht, 1977; Fersht and Dingwall, 1979). However, this contradicts the findings of Fournier et al. (2011), whose bioinformatic analyses would suggest the two-alphabet model be rejected on this cherry (IV). Our analysis, however, is more comprehensive because it considers alphabet expansion across the entire Class I and II aaRS trees, as opposed to just the IleRS-ValRS-LeuRS clade.

Third, these results suggest that acidic residues (Asp and Glu) either a) preceded their amide counterparts (Asn and Gln), or b) were once coded interchangeably with their respective amides in a stochastic fashion. Both scenarios are compatible with the observation that the naturally-occurring (Miller, 1953; Johnson et al., 2008) Asp and Glu are, respectively, metabolic precursors for Asn and Gln (Wong, 1975), and the role of pretranslational modification in encoding Gln and Asn in many living systems (Lamour et al., 1994; Hadd and Perona, 2014; Becker and Kern, 1998; Becker et al., 2000).

Fourth, the two-alphabet hypothesis was supported by the ingroup GlyRS-A (which forms a cherry with SerRS) but not the outgroup GlyRS-B (which forms a cherry with AlaRS). GlyRS-B, which is predominant in bacteria (Valencia-Sánchez et al., 2016), may have been a later innovation that outcompeted the GlyRS-A form mostly found in archaea and eukaryotes today. The ancestor of GlyRS-A and SerRS is therefore predicted to have been a promiscuous enzyme that recognised both amino acids, hence producing a stochastic genetic code.

Lastly, the Class II DK cherry was rejected. This is consistent with the hypothesis that Lys was already in the coding alphabet when the Class II LysRS spawned off AspRS, and consistent with with De Pouplana et al. (1998), who propose that convergently-evolved Class I and II LysRS were both present at the time of LUCA.

Conversely, some of our initial hypotheses were not supported by these experiments. In particular, we anticipated that the PT and FH aaRS cherries would be well-supported by the data, however these hypotheses were met with mixed support. When considering Class I and II aaRS individually, PT and FH were generally supported (Fig. 4), but when combined into a single analysis PT was rejected and FH was inconclusive (Table 2). This suggests that further information is required to link the two Class trees onto the same timescale. Alternatively, perhaps there were other (now-extinct) similar-functioned aaRS operating in parallel to the ancestor of ProRS and ThrRS, and the ancestor of HisRS and PheRS, i.e., a case of parafunctionalisation (Fournier et al., 2011). Furthermore, our negative control tests failed - two of our three “fake” cherries were supported by the aaRS data. This unexpected result suggests there are other evolutionary processes at play that cannot be explained by a two-alphabet aaRS model alone.

**Table 2:** Testing the two-alphabet hypothesis on both Classes of aaRS, using two different alignments. The posterior probability of each competing hypothesis is reported for each cherry *αβ*, where the total probability of a column sums to 1 (rounded to 2 decimal places). The leading hypothesis of the ancestral state is indicated. The last three cherries are negative controls and were expected to be rejected by this model.

| | aaRS cherry $\alpha\beta$ | | | | | | | | | Fake cherry $\alpha\beta$ | | |
| --- | --- | --- | --- | --- | --- | --- | --- | --- | --- | --- | --- | --- |
|  | WY | IV | EQ | DN | AG | SG | PT | FH | DK | LS | CF | NQ |
| Urzyme-I and II |  |  |  |  |  |  |  |  |  |  |  |  |
| $\alpha + \beta \rightarrow \alpha + \beta$ | 0.00 | 0.03 | 0.00 | 0.01 | **0.99** | **1.00** | **1.00** | **0.99** | **1.00** | **0.97** | 0.00 | 0.03 |
| $\alpha/\beta \rightarrow \alpha + \beta$ | 0.14 | **0.91** | **0.54** | **0.60** | 0.01 | 0.00 | 0.00 | 0.00 | 0.00 | 0.01 | 0.10 | **0.64** |
| $\alpha \rightarrow \alpha + \beta$ | 0.00 | 0.05 | 0.04 | 0.39 | 0.00 | 0.00 | 0.00 | 0.00 | 0.00 | 0.00 | **0.90** | 0.23 |
| $\beta \rightarrow \alpha + \beta$ | **0.86** | 0.00 | 0.42 | 0.00 | 0.00 | 0.00 | 0.00 | 0.01 | 0.00 | 0.02 | 0.00 | 0.10 |
| **Ancestral:** | Y | I/V | E/Q | D/N | A+G | S+G | P+T | F+H | D+K | L+S | C | N/Q |
| Cat-I and II |  |  |  |  |  |  |  |  |  |  |  |  |
| $\alpha + \beta \rightarrow \alpha + \beta$ | 0.00 | 0.00 | 0.04 | 0.00 | **1.00** | 0.00 | **0.90** | 0.49 | **0.99** | **0.67** | 0.00 | 0.00 |
| $\alpha/\beta \rightarrow \alpha + \beta$ | 0.00 | **0.99** | 0.39 | **0.76** | 0.00 | **0.90** | 0.06 | 0.06 | 0.00 | 0.00 | 0.02 | **0.79** |
| $\alpha \rightarrow \alpha + \beta$ | 0.00 | 0.01 | **0.57** | 0.24 | 0.00 | 0.10 | 0.04 | 0.00 | 0.00 | 0.00 | **0.98** | 0.21 |
| $\beta \rightarrow \alpha + \beta$ | **1.00** | 0.00 | 0.00 | 0.00 | 0.00 | 0.00 | 0.00 | **0.51** | 0.01 | 0.33 | 0.00 | 0.00 |
| **Ancestral:** | Y | I/V | E | D/N | A+G | S/G | P+T | H | D+K | L+S | C | N/Q |

### 4.2 Estimating the age of life and LUCA

Several previous studies have estimated the age of life and the age of LUCA through phylogenetic analysis of protein sequences. Their trees are often rooted early in Earth’s history; sometimes in the Hadean eon (4.0 - 4.6 Ga; Moody et al. (2024); Betts et al. (2018); Mahendrarajah et al. (2023)) and sometimes older than the Earth itself (Zhu et al., 2019). These studies make the usual phylogenetic assumption of life evolving at a steady pace in a gradualistic, temporally-regular fashion; at a clock rate that may have varied across lineages, but was constant through time on average (e.g., uncorrelated or autocorrelated clock models), and with a constant amino acid coding alphabet. If the aaRS did actually evolve in this manner, this would require that life came into existence at least 3 billion years before the Earth (Fig. 7A).

In reality, there are several evolutionary processes that make these assumptions difficult to justify, which will have consequences on the estimated date of LUCA and the origin of life. First, any pre- or post-LUCA refinements made to the coding table would have resulted in rapid, sweeping changes throughout the proteome. As we demonstrated, the ages of proteins generated under an expanding alphabet are overestimated when this process is unaccounted for (Fig. 7). Whereas, application of the resub model partially corrected this bias (that is, assuming that life did indeed originate on Earth and not extraterrestrially). Joining these aaRS bifurcation events, and others, into a multi-epoch multi-alphabet model would likely combine the effect we identified in individual cherries. Second, the initial forms of life probably evolved quite rapidly, due to low replicative fidelity and limited selective constraints; together leading to a higher clock rate than what we see in cellular systems today. Third, evolution often occurs in sudden punctuated bursts, where many changes occur at the time of speciation (i.e., on the tree nodes; Eldredge et al. (1972)). This is true not just for the early aaRS functional diversification events, but also other events after LUCA like the splitting of bacterial and archaeal lineages (Douglas et al., 2024b). Failing to account for this process can lead to an overestimation of divergence ages (Douglas et al., 2024b). While our work here does not proclaim a new and concrete date of abiogenesis, we have provided a step towards a better understanding of the complete timeline.

### 4.3 Two scenarios on the origin of coding

These findings offer further insight into competing theories concerning the origin of genetic coding, and therefore life as we know it (Kondratyeva et al., 2022). The prevailing *RNA World* hypothesis posits that protein coding originated in an environment governed by populations of self-reproducing RNA molecules, including ribozymes that carried out life’s chemical reactions (Janzen et al., 2020). In this scenario, protein synthesis was initially performed by ribozymal aminoacyl-tRNA synthetases (Wong et al., 2016). In contrast, the *Nucleopeptide World* proposes that coding originated in an environment governed by both RNA and peptides, with the proteinaceous aaRS serving a central integrating role between the two types of biomolecules (Carter Jr and Wills, 2021). Our observations in this study – that the Class I and II aaRS coevolved with each other and with the code itself – provide evidence in favour of this scenario over an RNA world.

Indeed, growing evidence supports a peptide-RNA origin of coding, as recently reviewed by Saad (2018); Kondratyeva et al. (2022); Wills and Carter Jr (2018). Complementary to our findings here, there are several lines of evidence attesting to this scenario. First, the near-universality of proteins/peptides that stabilise ribosomes, which implies that both types of biomolecule were involved at the dawn of ribosome-directed protein synthesis (Fox, 2010). Second, the observation that ribozymes have low complexities and limited catalytic abilities (Bernhardt, 2012); in particular the cryptic aaRS ribozymes, which have never been observed in nature, and to the best of our knowledge, have yet to be successfully engineered into synthetic ribozymes that perform both aaRS reactions – activation and charging (Lee et al., 2000; Illangasekare and Yarus, 1999; Niwa et al., 2009). Third, the capability of ancestral aaRS experimental models (urzymes) at performing both reactions necessary for tRNA aminoacylation at rates several orders of magnitude faster than uncatalysed (Li et al., 2011; Patra et al., 2024; Pham et al., 2010). These urzymes acylate ancestral tRNA models known as minihelices quite efficiently (Tang et al., 2024), and thrive even with the supposedly latecoming, but now-vital, histidine and lysine residues removed from the active site (Tang et al., 2023, 2024).

### 4.4 Conclusion and future outlook

We have opened up a more robust means to do phylogenetic inference on ancient proteins, while testing the hypothesis of alphabet expansion, all within a sound Bayesian framework. This in turn provides an important first step toward having up to twenty coding alphabet epochs informed by aaRS bifurcation events, paving the way for the study of ancestral protein quasispecies that were stochastically assembled from reduced alphabets. This article also offers further evidence in favour of a peptide-RNA scenario of life’s origins.

## 5 Limitations and assumptions

1. We considered just two epochs of coding alphabets at a time – the modern 20-amino-acid alphabet, and an ancestral 19-amino-acid alphabet. In reality, there were likely several epochs (18, 17, …). Our work here can be seamlessly extended to cover multiple epochs.
2. We considered the 20 standard amino acids – and not selenocysteine or pyrrolysine, or any other amino acids that may have once been in the coding alphabet.
3. We linked the two aaRS trees through temporal information (i.e., divergence dating). However it is known that aaRS did not evolve through clock-like processes. To address this issue, we used the gamma spike clock model, which assumed clock-like evolution along the lineages and abrupt evolution on the branch points (Douglas et al., 2024b).
4. Our model assumes just one ancestral genetic code at any given point in time, as opposed to multiple codes that operate across different organisms or antecedent protocells.
5. Standard phlyogenetic assumptions were made, including strictly bifurcating trees (i.e., vertical evolution), and independence between sites in the alignment.
6. The coding function of an ancestral aaRS might not be the same as its substrate specificity. For example, the ancestor of TrpRS an TyrRS might have been a promiscuous enzyme that recognised both amino acids, however if tryptophan was not abundant, then it would be absent from the coding alphabet.

## 6 Data, materials, and software availability

Our GPL-licensed source code is available as the resub package for BEAST 2. The software package, including the datasets and XML file configurations used in this article, are available at https://github.com/jordandouglas/resub.

## 7 Acknowledgements

The authors gratefully thank David Bryant for his early help in designing an algorithm for reducing amino acid substitution matrices. This work was supported by the Alfred P. Sloan Foundation Matter-to-Life program Grant number G-2021-16944. J.D. received additional research support from the MBIE (New Zealand Ministry of Business, Innovation & Employment) Beyond Prediction Data Science Research Programme (grant number UOAX1932). The authors wish to acknowledge the use of New Zealand eScience Infrastructure (NeSI) high performance computing facilities. New Zealand’s national facilities are provided by NeSI and funded jointly by NeSI’s collaborator institutions and through the Ministry of Business, Innovation & Employment’s Research Infrastructure programme. URL https://www.nesi.org.nz.

