## Supporting Information for "Reduced amino acid substitution matrices find traces of ancient coding alphabets in modern day proteins"

February 19, 2025

### 1 Simulation studies

#### 1.1 Coverage simulation studies

We performed coverage simulation studies[1] to validate our phylogenetic model. This involved the following steps, across 100 replicates. First, we simulated a set of binary rooted time-trees $\mathcal{T}$ as birth-death processes, with birth rate $\lambda$ and death rate $\mu$ sampled from their prior distributions, with extinct lineages removed. These trees were simulated such that the average tree height was approximately 1 – 2 substitutions per site (depending on the number of taxa) to achieve mutational saturation. Second, we selected the substitution model indicator $\mathbb{I}_s \in \{0, 1, 2, 3\}$. This step was not stochastic - rather we ensured that $\mathbb{I}_s = 0$ on 50/100 replicates, $\mathbb{I}_s = 1$ on 25/100, and $\mathbb{I}_s = 2$ on 13/100, and $\mathbb{I}_s = 3$ on 12/100. Third, we uniformly at random sampled a cherry $\alpha\beta \in \{$AG, DN, DK, EQ, FH, IV, PT, SG, WY$\}$. Fourth, we simulated amino acid sequences down the tree $\mathcal{T}$ using the `resub(`$\alpha\beta$`)` model, including model indicator $\mathbb{I}_s$. We used the aaRS substitution model (Table S1 and Fig. S4) and gamma spike model[2] during this step. Fifth, we performed MCMC on each simulated dataset to estimate the tree and its parameters. Lastly, we compared the known values of parameters with their posterior estimates.

We validated this approach using coverage simulation studies with $N = 20$ (Fig. S1) and $N = 40$ (Fig. S2) taxa. These experiments confirmed the true value of a parameter lies in its 95% credible interval approximately 95% of the time; thereby providing confidence in the correctness of our method, and its ability to recover parameter estimates from data simulated under a known model. Under these conditions, not all parameters were informed by the data however. For example, the $N = 20$ trees provided no information about the spike gamma distribution shape $S_\alpha$.

During these experiments, the following priors were used:

- Birth rate
  - $\lambda \sim \text{LogNormal}(\text{mean} = 2, \sigma = 0.2)$ when $N = 20$
  - $\lambda \sim \text{LogNormal}(\text{mean} = 3, \sigma = 0.2)$ when $N = 40$
- Reproduction number $\frac{\lambda}{\mu} - 1 \sim \text{Exponential}(\text{mean} = 5)$
- Gradual relaxed clock standard deviation $\sim \text{Gamma}(\alpha = 5, \beta = 0.05)$
- Spike mean $S_\mu \sim \text{LogNormal}(\text{mean} = 0.01, \sigma = 1.2)$
- Spike shape $S_\alpha \sim \text{LogNormal}(\text{mean} = 2, \sigma = 0.5)$
- Gamma rate heterogeneity shape $\sim \text{Exponential}(\mu = 1)$
- Transition proportion $\nu \sim \text{Beta}(\alpha = 5, \beta = 5)$
- Relative transition age $\frac{t_e}{t_h} \sim \text{Beta}(\alpha = 6, \beta = 2)$ where $t_h$ is the root height
- Amino acid equilibrium frequencies $\pi \sim \text{Dirichlet}(\alpha_1 = 4, \alpha_2 = 4, \dots, \alpha_{20} = 4)$ - and fixed to the aaRS empirical substitution model during simulation (Table S1).
- Amino acid exchangeability relative rates $\mathbf{r} \sim \text{LogNormal}(\text{mean} = 1, \sigma = 1)$ - and fixed to the aaRS empirical substitution model during simulation (Fig. S4).

- Model indicator $\mathbb{I}_s = \begin{cases} 0 & \text{w.p. } \frac{1}{2} \\ 1 & \text{w.p. } \frac{1}{4} \\ 2 & \text{w.p. } \frac{1}{8} \\ 3 & \text{w.p. } \frac{1}{8} \end{cases}$

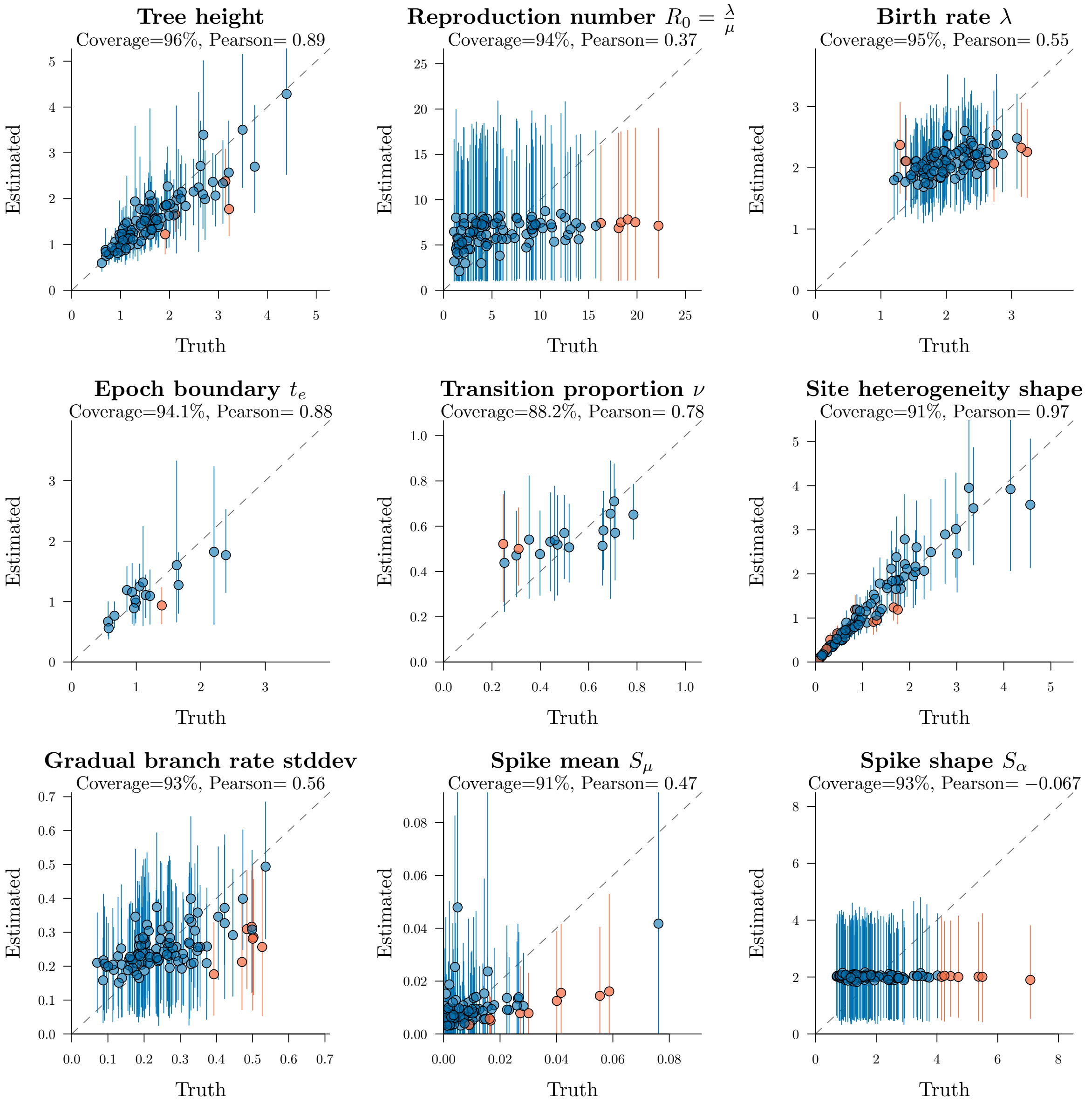

Fig. S1: Well-calibrated simulation study on trees with $N = 20$ taxa and $L = 100$ amino acid sites under the `resub` model. 100 replicates were performed in total, with $t_e$ and $\nu$ plots being conditional on the true model indicator $\mathbb{I}_s$ being estimated as the leading hypothesis. Points are coloured blue if the true value is within the 95% credible interval, and orange otherwise. The coverage is the proportion of replicates whose true value is within the 95% credible interval (close to 95% indicating good coverage). The Pearson correlation between true values and mean estimates is also reported.

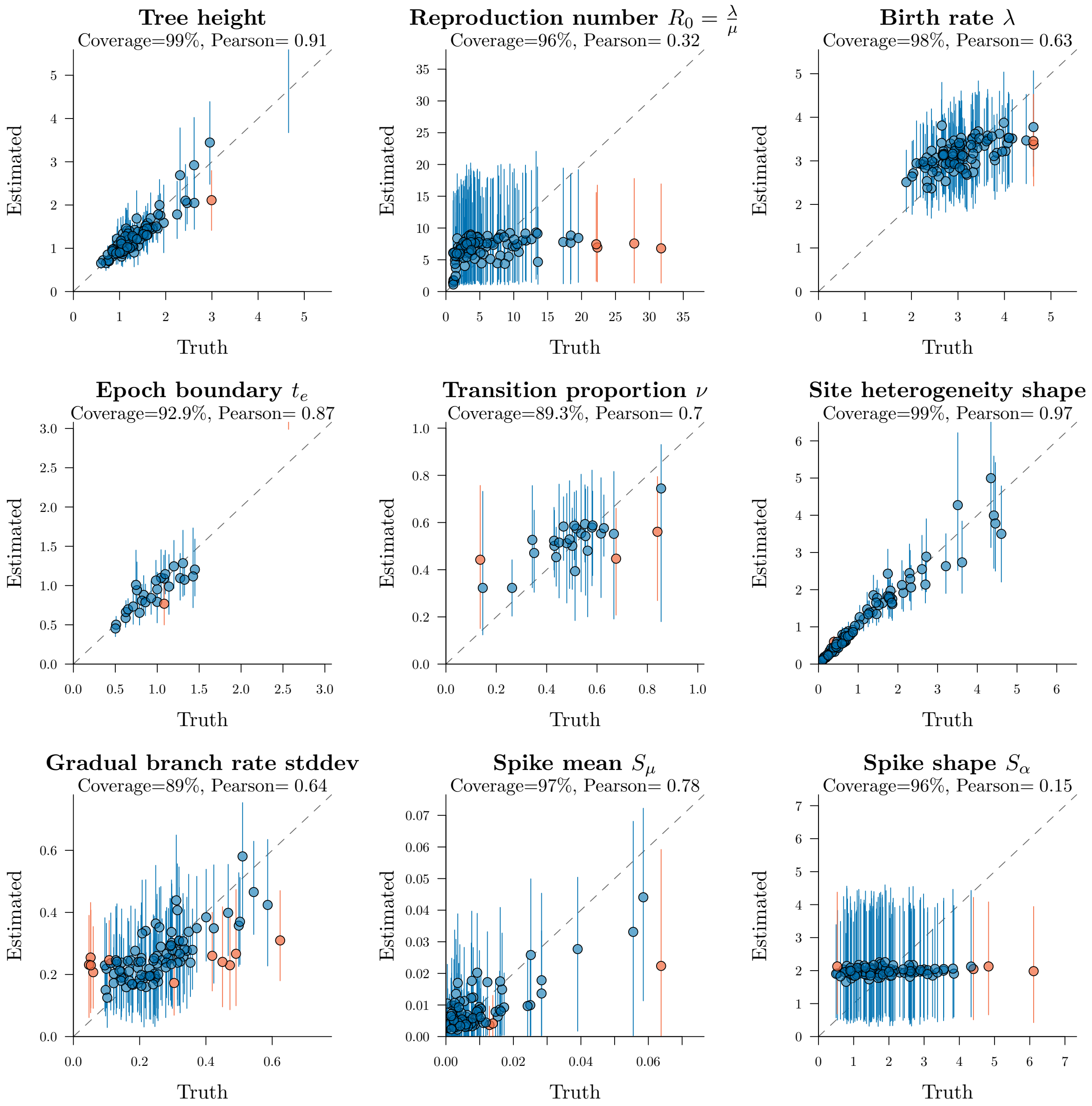

Fig. S2: Well-calibrated simulation study on trees with $N = 40$ taxa and $L = 100$ amino acid sites. Refer to Fig. S1 for figure notation.

#### 1.2 Bayesian model averaging

By performing additional simulation studies, we characterised how accurately our new method can identify the true substitution model $\mathbb{I}_s$ on data simulated under a known sibstituiton model. The simulations were conducted under the same conditions described in the section above, for varying alignment lengths $L = (100, 200, 500)$ and taxon counts $N = (20, 40)$. As shown in Fig 3 of the main article, this method can usually identify whether `resub` is appropriate, with confidence increasing with the size of the tree, and very rarely wrong. However, as shown in Fig. S3, it is much more difficult, but still possible, to discriminate between the various cases of resub without large datasets (i.e., it is challenging to discriminate between $\mathbb{I}_s = 1, 2,$ and 3). These experiments confirm this approach is statistically consistent, with accuracy improving with both longer alignments and larger trees.

We compared varying hypotheses using Bayes factors. Following the guidelines of Kass and Raftery,[3] a Bayes factor $B_h$ of 10 indicates "strong" support in favour of hypothesis $h$. This threshold corresponds to the following posterior probabilities:

$$B_h = \frac{p(\mathbb{I}_s = h|D)}{p(\mathbb{I}_s = h)} \div \frac{p(\mathbb{I}_s \neq h|D)}{p(\mathbb{I}_s \neq h)} \tag{1}$$

$$\begin{aligned} \Rightarrow p(\mathbb{I}_s = 0|D) &> 0.91 \text{ when } B_0 > 10. \\ p(\mathbb{I}_s = 1|D) &> 0.77 \text{ when } B_1 > 10. \\ p(\mathbb{I}_s = 2|D) &> 0.59 \text{ when } B_2 > 10. \\ p(\mathbb{I}_s = 3|D) &> 0.59 \text{ when } B_3 > 10. \end{aligned} \tag{2}$$

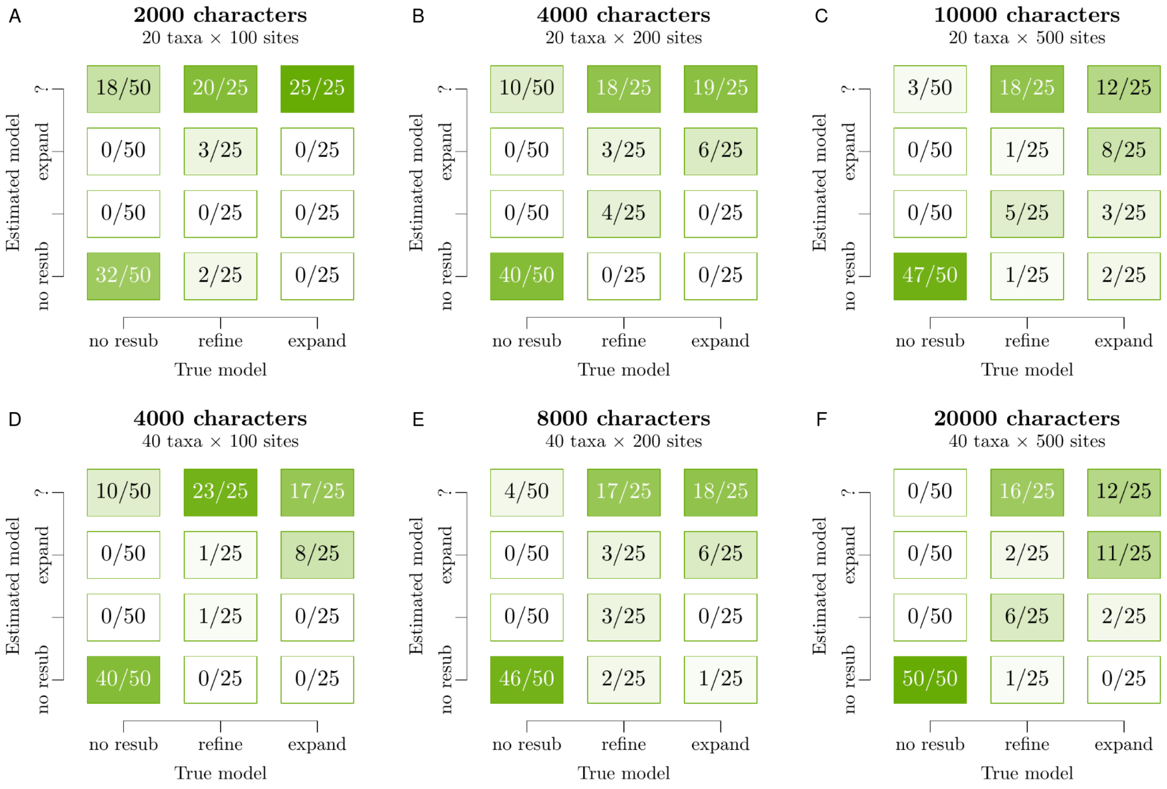

Fig. S3: Bayesian model averaging on simulated data of varying sizes. 100 datasets were simulated and 100 MCMC chains run for each panel in the figure, and the inferred model was classified into no resub, refinement, expansion, or unclassified using a Bayes factor threshold of 10. These results demonstrate that, while the method is rarely wrong, it is still challenging to discriminate between refinement and expansion. This limitation improves with larger trees and longer alignments. Note that this figure is an alternative view of the same data presented in Fig. 3 of the main article.

#### 1.3 Estimating tree height bias on simulated data

In Fig. 7 of main article, we assessed the impact of applying either model ($\mathbb{I}_s$ fixed at 0, or fixed at its true value $> 0$) to data generated under the latter. We showed that the former produced biased estimates of tree height and the age of the old epoch. These trees were calibrated time trees, meaning that the node ages represented time units informed by the presence of sampled ancestral nodes on the tree. Sampled ancestors were simulated under a birth-death-sampling model[4] with the sampling proportion fixed to 0.3. The simulated alignments consisted of $L = 200$ amino acid sites, $N = 40$ extant taxa, and a variable number of non-extant taxa (mean: 24, 95% HPD: (1, 65), range: (1, 146)). Data were simulated and inferred under a relaxed clock model.[5] The following priors were used in this experiment:

- Clock rate
  - $\mu_C = 1$ during simulation
  - $\mu_C \sim \text{LogNormal}(\text{mean} = 1, \sigma = 1)$ during inference
- Birth rate
  - $\lambda \sim \text{LogNormal}(\text{mean} = 2, \sigma = 0.2)$ during simulation
  - $\lambda \sim \text{LogNormal}(\text{mean} = 10, \sigma = 2)$ during inference (i.e., an uninformed prior)
- Reproduction number $\frac{\lambda}{\mu} - 1 \sim \text{Exponential}(\text{mean} = 1)$
- Gradual relaxed clock standard deviation $\sim \text{Gamma}(\alpha = 5, \beta = 0.05)$
- Gamma rate heterogeneity shape $\sim \text{Exponential}(\mu = 1)$
- Transition proportion $\nu \sim \text{Beta}(\alpha = 5, \beta = 5)$
- Relative transition age
  - $\frac{t_e}{t_h} \sim \text{Beta}(\alpha = 10, \beta = 2)$ during simulation
  - $\frac{t_e}{t_h}$ fixed at true value during inference
- Model indicator
  - $\mathbb{I}_s = \begin{cases} 0 & \text{w.p. } 0 \\ 1 & \text{w.p. } \frac{1}{2} \\ 2 & \text{w.p. } \frac{1}{4} \\ 3 & \text{w.p. } \frac{1}{4} \end{cases}$ during simulation
  - $\mathbb{I}_s$ fixed at either 0 or the true value during inference
- Amino acid equilibrium frequencies $\sim \text{Dirichlet}(\alpha_A = 4, \alpha_C = 4, \dots, \alpha_Y = 4)$
- Amino acid exchangeability relative rates fixed to the aaRS empirical substitution model.

Under this prior distribution, the true tree height $t_h$ averaged 3.1 substitutions per site (95% credible interval: (1.1, 5.8)). However, when doing inference, we gave $\lambda$ an uninformed prior, which resulted in the tree height averaging 34 (0.009, 126) substitutions per site. As a result, the tree height during inference was informed by the data, and not by the prior.

#### 2 An empirical substitution model of the aaRS

We estimated the exchangeability matrix **r** from a joint Bayesian analysis of the Class I and II aaRS catalytic domains. This matrix was used for simulating data in our well-calibrated simulation studies. This matrix is shown in Fig. S4. The estimated frequencies are shown in Table S1.

| **Amino acid** | **Estimated frequency** |
|---|---|
| A | 0.07611 |
| C | 0.01206 |
| D | 0.04293 |
| E | 0.07649 |
| F | 0.05826 |
| G | 0.06448 |
| H | 0.02521 |
| I | 0.0684 |
| K | 0.05763 |
| L | 0.09824 |
| M | 0.02922 |
| N | 0.03194 |
| P | 0.03257 |
| Q | 0.03192 |
| R | 0.06107 |
| S | 0.0547 |
| T | 0.04757 |
| V | 0.06342 |
| W | 0.02267 |
| Y | 0.04511 |

Table S1: Estimated amino acid frequencies in the Class I and II aaRS catalytic domains.

**Amino acid exchangability rates in the aaRS**

| | A | R | N | D | C | Q | E | G | H | I | L | K | M | F | P | S | T | W | Y | V |
|---|---|---|---|---|---|---|---|---|---|---|---|---|---|---|---|---|---|---|---|---|
| A | | | | | | | | | | | | | | | | | | | | |
| R | 0.39 | | | | | | | | | | | | | | | | | | | |
| N | 0.34 | 1.5 | | | | | | | | | | | | | | | | | | |
| D | 0.38 | 0.22 | 4.7 | | | | | | | | | | | | | | | | | |
| C | 4.4 | 0.55 | 2.4 | 0.28 | | | | | | | | | | | | | | | | |
| Q | 0.64 | 1.4 | 2.4 | 0.96 | 0.31 | | | | | | | | | | | | | | | |
| E | 0.56 | 0.49 | 1.3 | 3.6 | 0.14 | 3 | | | | | | | | | | | | | | |
| G | 2 | 0.072 | 1.6 | 0.93 | 0.72 | 0.16 | 0.16 | | | | | | | | | | | | | |
| H | 0.47 | 0.9 | 4.6 | 0.59 | 0.59 | 2.9 | 0.48 | 0.28 | | | | | | | | | | | | |
| I | 0.27 | 0.19 | 0.19 | 0 | 0.51 | 0.36 | 0 | 0 | 0.28 | | | | | | | | | | | |
| L | 0.32 | 0.21 | 0.15 | 0 | 1.8 | 0.53 | 0.11 | 0.033 | 0.3 | 3.9 | | | | | | | | | | |
| K | 0.45 | 4.4 | 2.1 | 0.86 | 0.12 | 3.5 | 1.4 | 0.44 | 0.7 | 0.13 | 0.28 | | | | | | | | | |
| M | 1.2 | 0.36 | 0.2 | 0 | 1.2 | 2.4 | 0.09 | 0.1 | 0.48 | 3.5 | 5.4 | 0.49 | | | | | | | | |
| F | 0.34 | 0.079 | 0 | 0 | 0.94 | 0 | 0.12 | 0 | 0.9 | 1 | 1.7 | 0 | 1.9 | | | | | | | |
| P | 0.87 | 0.3 | 0.66 | 0.44 | 0.57 | 0.39 | 0.33 | 0.27 | 0.27 | 0.15 | 0.2 | 0.73 | 0.27 | 0.074 | | | | | | |
| S | 4 | 0.8 | 3.2 | 1.3 | 3.3 | 1 | 0.61 | 1.4 | 1.7 | 0 | 0.11 | 1.3 | 0.68 | 0.11 | 1 | | | | | |
| T | 1.7 | 0.52 | 1.6 | 0.57 | 2.9 | 1.2 | 0.57 | 0.35 | 1.3 | 0.76 | 0.52 | 0.81 | 1.5 | 0.16 | 0.78 | 5.1 | | | | |
| W | 0.15 | 0.37 | 0.098 | 0.24 | 0.55 | 0.41 | 0 | 0.099 | 0.84 | 0.097 | 0.38 | 0 | 0.74 | 2 | 0.094 | 0.37 | 0.44 | | | |
| Y | 0.56 | 0.28 | 1.1 | 0.15 | 0.77 | 0.38 | 0.14 | 0.066 | 3.8 | 0.24 | 0.48 | 0.44 | 0.76 | 5.9 | 0.26 | 0.23 | 0.5 | 2.1 | | |
| V | 2.3 | 0.15 | 0.34 | 0.14 | 4.4 | 0.34 | 0.24 | 0 | 0.39 | 8.7 | 1.8 | 0.23 | 1.7 | 0.73 | 0.45 | 0.42 | 2.9 | 0.24 | 0.55 | |

Fig. S4: Amino acid exchangeability matrix estimated from the Class I and II aaRS catalytic domains. The numbers shown are relative transition rates, with reds indicating faster rates, and blue indicating slow rates. The 0 rates correspond to parameters that were deemed as unnecessary by stochastic variable selection. The values shown here are median posterior estimates.

### 3 Biological datasets

#### 3.1 Prior distributions

The following prior distributions were used to generate the resub screening results presented in Fig. 4 of the main article:

- Birth rate $\lambda \sim \text{LogNormal}(\text{mean} = 1, \sigma = 2)$
- Reproduction number $\frac{\lambda}{\mu} - 1 \sim \text{Exponential}(\text{mean} = 5)$
- Gradual relaxed clock standard deviation $\sim \text{Gamma}(\alpha = 5, \beta = 0.05)$
- Spike mean $S_\mu \sim \text{LogNormal}(\text{mean} = 0.01, \sigma = 1.2)$
- Spike shape $S_\alpha \sim \text{LogNormal}(\text{mean} = 2, \sigma = 0.5)$
- Gamma rate heterogeneity shape $\sim \text{Exponential}(\mu = 1)$
- Amino acid equilibrium frequencies $\sim \text{Dirichlet}(\alpha_A = 4, \alpha_C = 4, \ldots, \alpha_Y = 4)$
- Amino acid exchangeability rates $\mathbf{r} \sim \text{LogNormal}(\text{mean} = 1, \sigma = 1)$
- Transition proportion $\nu \sim \text{Beta}(\alpha = 4, \beta = 4)$
- Relative transition age $\frac{t_e}{t_h} \sim \text{Beta}(\alpha = 6, \beta = 2)$
- Model indicator $\mathbb{I}_s = \begin{cases} 0 & \text{w.p. } \frac{1}{2} \\ 1 & \text{w.p. } \frac{1}{4} \\ 2 & \text{w.p. } \frac{1}{8} \\ 3 & \text{w.p. } \frac{1}{8} \end{cases}$

#### 3.2 Site compositions

To better understand these biological datasets, we counted the number of alignment sites that contain both members of each cherry (Table S2) and the empirical frequency of each amino acid (Table S3). These observations confirm that each amino acid and each cherry were well-represented across the datasets.

| Dataset | AG | SG | IV | PT | DK | DN | EQ | FH | WY | LS | NQ | CF |
|---|---|---|---|---|---|---|---|---|---|---|---|---|
| Transferrin | 65 | 75 | 74 | 36 | 70 | 74 | 83 | 8 | 7 | 55 | 49 | 7 |
| Papain-like protease | 71 | 76 | 75 | 55 | 75 | 79 | 72 | 25 | 12 | 76 | 61 | 9 |
| Trypsin | 135 | 151 | 114 | 110 | 108 | 101 | 109 | 55 | 34 | 176 | 90 | 35 |
| Carbonic anhydrase | 125 | 129 | 109 | 78 | 94 | 96 | 108 | 38 | 14 | 142 | 79 | 14 |
| Elongation factors | 125 | 90 | 229 | 65 | 155 | 99 | 123 | 22 | 12 | 120 | 63 | 12 |
| I3G phosphate synthase | 49 | 39 | 77 | 17 | 56 | 42 | 58 | 8 | 4 | 42 | 31 | 7 |
| DNA repair | 113 | 130 | 172 | 90 | 185 | 150 | 170 | 47 | 24 | 215 | 115 | 29 |
| Cytochrome P450 | 151 | 124 | 117 | 82 | 103 | 88 | 121 | 45 | 21 | 112 | 71 | 18 |
| Glutathione S-transferase | 68 | 67 | 82 | 39 | 63 | 56 | 66 | 43 | 16 | 69 | 52 | 19 |
| Triosephosphate isomerase | 99 | 97 | 112 | 49 | 81 | 88 | 87 | 41 | 10 | 94 | 89 | 24 |
| Class I urzyme bacterial | 71 | 81 | 90 | 49 | 53 | 57 | 54 | 54 | 31 | 91 | 60 | 42 |
| Class I urzyme | 107 | 111 | 115 | 78 | 81 | 85 | 90 | 81 | 54 | 119 | 96 | 69 |
| Class I catalytic domain | 127 | 130 | 133 | 90 | 104 | 101 | 112 | 100 | 81 | 134 | 114 | 88 |
| Class II urzyme bacterial | 95 | 89 | 97 | 63 | 73 | 81 | 95 | 63 | 42 | 106 | 90 | 43 |
| Class II urzyme | 123 | 126 | 128 | 98 | 103 | 118 | 125 | 103 | 75 | 134 | 123 | 76 |
| Class II catalytic domain | 133 | 132 | 132 | 94 | 96 | 113 | 108 | 107 | 92 | 136 | 109 | 88 |

Table S2: The number of alignment sites that contain both members of each cherry.

| **Dataset** | **A** | **C** | **D** | **E** | **F** | **G** | **H** | **I** | **K** | **L** | **M** | **N** | **P** | **Q** | **R** | **S** | **T** | **V** | **W** | **Y** |
|---|---|---|---|---|---|---|---|---|---|---|---|---|---|---|---|---|---|---|---|---|
| Transferrin | 0.0904 | 0.047 | 0.0618 | 0.0586 | 0.0367 | 0.0789 | 0.0206 | 0.0368 | 0.0754 | 0.0823 | 0.014 | 0.0382 | 0.0435 | 0.0358 | 0.0375 | 0.0774 | 0.0525 | 0.0646 | 0.0127 | 0.0351 |
| Papain-like protease | 0.0759 | 0.0342 | 0.051 | 0.0587 | 0.0332 | 0.0978 | 0.0218 | 0.0538 | 0.059 | 0.0566 | 0.016 | 0.0545 | 0.0408 | 0.0371 | 0.0386 | 0.0775 | 0.0527 | 0.0677 | 0.0238 | 0.0494 |
| Trypsin | 0.0674 | 0.0447 | 0.0478 | 0.0446 | 0.0252 | 0.0916 | 0.0308 | 0.0464 | 0.042 | 0.0882 | 0.0159 | 0.0412 | 0.0597 | 0.042 | 0.0474 | 0.0785 | 0.0535 | 0.0753 | 0.0241 | 0.0336 |
| Carbonic anhydrase | 0.06 | 0.0121 | 0.0515 | 0.0653 | 0.0377 | 0.0664 | 0.0359 | 0.0439 | 0.0469 | 0.0959 | 0.018 | 0.0469 | 0.0643 | 0.0455 | 0.0424 | 0.0878 | 0.0603 | 0.0664 | 0.0172 | 0.0355 |
| Elongation factors | 0.0721 | 0.011 | 0.0539 | 0.0839 | 0.0299 | 0.0807 | 0.0212 | 0.0825 | 0.0866 | 0.0776 | 0.0206 | 0.0341 | 0.0518 | 0.0282 | 0.0494 | 0.0439 | 0.0547 | 0.0882 | 0.00765 | 0.0221 |
| I3G phosphate synthase | 0.1 | 0.0079 | 0.0578 | 0.0933 | 0.032 | 0.0569 | 0.0106 | 0.0787 | 0.0663 | 0.111 | 0.0164 | 0.0324 | 0.0386 | 0.0295 | 0.0646 | 0.0634 | 0.0355 | 0.0769 | 0.00332 | 0.0247 |
| DNA repair | 0.0686 | 0.00872 | 0.0663 | 0.0864 | 0.0384 | 0.0567 | 0.0146 | 0.063 | 0.084 | 0.102 | 0.0201 | 0.0428 | 0.0451 | 0.0422 | 0.0542 | 0.0683 | 0.0458 | 0.0553 | 0.00808 | 0.0296 |
| Cytochrome P450 | 0.0958 | 0.0106 | 0.0669 | 0.0726 | 0.0447 | 0.0622 | 0.0284 | 0.0483 | 0.0338 | 0.115 | 0.0246 | 0.0257 | 0.0625 | 0.0343 | 0.0789 | 0.0483 | 0.0524 | 0.0658 | 0.00841 | 0.0203 |
| Glutathione S-transferase | 0.0809 | 0.00726 | 0.0546 | 0.0799 | 0.05 | 0.0553 | 0.0218 | 0.0536 | 0.0816 | 0.115 | 0.0284 | 0.0354 | 0.0485 | 0.0367 | 0.0467 | 0.0469 | 0.0376 | 0.0657 | 0.0118 | 0.042 |
| Triosephosphate isomerase | 0.108 | 0.0154 | 0.0433 | 0.073 | 0.0331 | 0.0811 | 0.0229 | 0.0815 | 0.0591 | 0.0849 | 0.0186 | 0.0474 | 0.0316 | 0.0422 | 0.0378 | 0.0617 | 0.0471 | 0.076 | 0.0114 | 0.0242 |
| Class I urzyme bacterial | 0.0646 | 0.0105 | 0.0592 | 0.0371 | 0.053 | 0.087 | 0.0436 | 0.0643 | 0.0527 | 0.0914 | 0.0286 | 0.0393 | 0.05 | 0.028 | 0.0499 | 0.0558 | 0.0531 | 0.0666 | 0.0175 | 0.0476 |
| Class I urzyme | 0.0654 | 0.0113 | 0.057 | 0.0386 | 0.0541 | 0.0823 | 0.0442 | 0.0654 | 0.0549 | 0.091 | 0.0286 | 0.0377 | 0.0498 | 0.0275 | 0.05 | 0.0584 | 0.0525 | 0.0666 | 0.0181 | 0.0468 |
| Class I catalytic domain | 0.0687 | 0.0131 | 0.0504 | 0.0524 | 0.0581 | 0.06 | 0.0406 | 0.0673 | 0.0616 | 0.09 | 0.0288 | 0.0346 | 0.0421 | 0.0324 | 0.0557 | 0.0579 | 0.0441 | 0.0638 | 0.0231 | 0.0551 |
| Class II urzyme bacterial | 0.0693 | 0.01 | 0.0467 | 0.086 | 0.0633 | 0.0592 | 0.03 | 0.0558 | 0.0468 | 0.0979 | 0.0308 | 0.0344 | 0.0441 | 0.0407 | 0.0698 | 0.0471 | 0.0547 | 0.0585 | 0.0127 | 0.0422 |
| Class II urzyme | 0.0647 | 0.0119 | 0.0467 | 0.0859 | 0.0655 | 0.0553 | 0.0302 | 0.0584 | 0.0495 | 0.0997 | 0.0312 | 0.0349 | 0.0418 | 0.0388 | 0.0672 | 0.0511 | 0.0534 | 0.0591 | 0.0128 | 0.0419 |
| Class II catalytic domain | 0.0629 | 0.0155 | 0.0407 | 0.0828 | 0.0636 | 0.0698 | 0.0256 | 0.0698 | 0.0422 | 0.103 | 0.0338 | 0.0312 | 0.0363 | 0.0348 | 0.0619 | 0.0501 | 0.0516 | 0.0668 | 0.015 | 0.0426 |

Table S3: Empirical amino acid frequencies in each dataset (rounded to 3 sf).

### 4 Joint aaRS phylogenetic analysis

#### 4.1 Prior distributions for calibrated root analysis

The following prior distributions were used in our Class I and II aaRS catalytic domain joint analyses, presented in Fig. 5 and 6 of the main article. This model featured a birth-death tree prior and a gamma spike clock model.

- Birth rates; one per Class $\lambda \sim \text{LogNormal}(\text{mean} = 1, \sigma = 1)$
- Reproduction numbers; one per Class $\frac{\lambda}{\mu} - 1 \sim \text{Exponential}(\text{mean} = 5)$
- Gradual relaxed clock standard deviations; one per Class $\sim \text{Gamma}(\alpha = 5, \beta = 0.05)$
- Spike means; one per Class $S_\mu \sim \text{LogNormal}(\text{mean} = 0.01, \sigma = 1.2)$
- Spike shapes; one per Class $S_\alpha \sim \text{LogNormal}(\text{mean} = 2, \sigma = 0.5)$
- Clock rate; amino acid substitutions per site per billion years $\sim \text{LogNormal}(\text{mean} = 0.1, \sigma = 1)$
- Relative mutation rates; one per Class $\sim \text{LogNormal}(\text{mean} = 1, \sigma = 1)$
- Gamma rate heterogeneity shapes; one per Class $\sim \text{LogNormal}(\text{mean} = 2, \sigma = 0.5)$
- Amino acid equilibrium frequencies $\sim \text{Dirichlet}(\alpha_A = 4, \alpha_C = 4, \ldots, \alpha_Y = 4)$
- Amino acid exchangeability rates $\mathbf{r} \sim \text{LogNormal}(\text{mean} = 1, \sigma = 1)$
- Transition proportion $\nu \sim \text{Beta}(\alpha = 5, \beta = 5)$
- Relative transition age
  - $t_e - t_a \sim \text{Laplace}(\lambda = 100)$ for most cherries, where $t_a$ is the estimated height of the cherry aaRS ancestor.
  - $\frac{t_e - t_a}{t_h} \sim \text{Beta}(\alpha = 2, \beta = 6)$ for the pre-translational modification cherries (EQ and DN).
- Model indicator $\mathbb{I}_s = \begin{cases} 0 & \text{w.p. } \frac{1}{2} \\ 1 & \text{w.p. } \frac{1}{4} \\ 2 & \text{w.p. } \frac{1}{8} \\ 3 & \text{w.p. } \frac{1}{8} \end{cases}$

Under this model, the overall evolutionary rate of a Class tree is equal to the clock rate times the relative mutation rate of that tree.

The transition boundary $t_e$ was *a priori* constrained to sensible estimates informed by the aaRS phylogeny. In the case of EQ and DN, we viewed the aaRS ancestor (i.e., the ancestor of GluRS and GlnRS; or AspRS and AsnRS) as being a lower bound on the coding alphabet bifurcation event, given that we already know that these ancestors were non-discriminating aaRS that behaved deterministically with a full alphabet (GlxRS and AsxRS). Therefore, $t_e$ was constrained between the time of this node $t_a$ and the tree root height $t_h$. In the general case, $t_e$ was strongly constrained to occur near the same height $t_a$ as the respective aaRS node. In the case of real cherries, this is straightforward. For example, in the WY cherry, $t_a$ is the time of the ancestor of TrpRS and TyrRS. In the case of the fake cherries, $t_a$ was linked to one of the other cherry ancestors, chosen arbitrarily. For LS, $t_a$ was the time of the SerRS/GlyRS ancestor; for CF, $t_a$ was linked to TrpRS/TyrRS; and for NQ, $t_a$ was linked to GluRS/GlnRS.

#### 4.2 Prior distributions for uncalibrated root analysis

In the aaRS analysis of Fig. 7 of the main article, we removed the root age divergence time prior, but retained the other calibration priors (LUCA, LBCA, LACA, LMCA, and LECA). We applied a relaxed clock[5] and a birth-death-skyline tree prior.[6] This skyline model allowed the speciation rate to vary at the top and bottom of the tree, by assuming two branching process epochs each with an i.i.d. net diversification rate. The epoch boundary was fixed at 4.2 Ga, approximately the age of LUCA. The priors of this skyline model were selected such that the root height was *a priori* centered around a 95% credible interval of 4.46 – 5.70 Ga. We used resub (with the WY, IV, EQ, DN, and SG cherries respetively) and the Null model ($\mathbb{I}_s = 0$), each with the following priors:

- Post-LUCA net diversification rates; one per Class $\lambda - \mu \sim \text{LogNormal}(\text{mean} = 10, \sigma = 1)$
- Pre-LUCA net diversification rates; one per Class $\lambda - \mu \sim \text{LogNormal}(\text{mean} = 3, \sigma = 0.05)$
- Reproduction numbers; one per Class $\frac{\lambda}{\mu} = 2$
- Sampling proportion; one per Class, fixed at $10^{-8}$ due to absence of sampled ancestors
- Gradual relaxed clock standard deviations; one per Class $\sim \text{Gamma}(\alpha = 5, \beta = 0.05)$
- Clock rate; amino acid substitutions per site per billion years $\sim \text{LogNormal}(\text{mean} = 0.1, \sigma = 1)$
- Relative mutation rates; one per Class $\sim \text{LogNormal}(\text{mean} = 1, \sigma = 1)$
- Gamma rate heterogeneity shapes; one per Class $\sim \text{LogNormal}(\text{mean} = 2, \sigma = 0.5)$
- Amino acid equilibrium frequencies $\sim \text{Dirichlet}(\alpha_A = 4, \alpha_C = 4, \dots, \alpha_Y = 4)$
- Amino acid exchangeability rates $\mathbf{r} \sim \text{LogNormal}(\text{mean} = 1, \sigma = 1)$
- Transition proportion $\nu \sim \text{Beta}(\alpha = 5, \beta = 5)$; parameter is not being used in the Null model
- Relative transition age $t_e - t_a \sim \text{Laplace}(\lambda = 20)$, where $t_a$ is the estimated height of the cherry aaRS ancestor; parameter is not being used in the Null model
- Model indicator $\mathbb{I}_s = \begin{cases} 0 & \text{w.p. } 0 \\ 1 & \text{w.p. } \frac{1}{2} \\ 2 & \text{w.p. } \frac{1}{4} \\ 3 & \text{w.p. } \frac{1}{4} \end{cases}$ when using resub, or fixed to 0 in the Null model
